# Eyes wide open: Frogs with reduction of auditory communication ability show lower eye, but higher corneal, investment

**DOI:** 10.1101/2025.11.07.687275

**Authors:** Jhael A. Ortega, Kate N. Thomas, Santiago R. Ron, Rayna C. Bell, Matthew K. Fujita, David J. Gower, Jeffrey W. Streicher, Ryan K. Schott

## Abstract

Vocal communication is the primary mechanism of mate attraction in frogs, but some species have lost the auditory structures needed to hear airborne sounds, raising the question of how they communicate in their absence. We hypothesized that frog species with auditory system reductions have compensated through enhanced visual ability, including through enlarged eyes. To test this trade-off hypothesis, we estimated relative eye and corneal investment across 264 species and tested for correlations with auditory reduction. Contrary to our expectations, we found that species with auditory reductions had smaller eye size and lower eye investment. Instead, we found that non-fossorial species with auditory reductions had significantly greater corneal investment, after controlling for habitat-based variation. Enlarged corneas enhance light sensitivity but are expected to have a lower metabolic cost than larger eyes. Relatively larger corneas may, in combination with other changes, allow frogs with auditory reductions to rely more on vision for communication, while at the same time minimizing energetic costs in small animals that are already metabolically constrained. Our findings suggest that the visual system plays an adaptive role in response to the reduction of auditory communication and highlights the condition of sensory systems as an important factor shaping frog evolution.

## Introduction

Acoustic communication is widespread across animals, serving as a key mechanism for mediating social interactions and reproductive activities in diverse ecological contexts (1). In vertebrates, acoustic signals are particularly effective at transmitting information over long distances and through acoustically complex environments (2), making them central to coordinating social and reproductive behaviours. This broad importance of vocal signals sets the stage for understanding frogs and toads (Anura; here collectively referred to as frogs), where acoustic communication plays a central role in social and reproductive biology and is widely considered a major driver of their evolutionary diversification. Frogs rely predominantly on vocal signals to mediate a range of behaviours, especially those related to courtship, mating, and territory defence (3). These vocalizations are diverse and context-dependent, broadly categorized into reproductive, aggressive, and defensive calls, each encompassing multiple subtypes (4,5). Advertisement calls, in particular, are the most widespread and taxonomically informative, typically produced by males to attract females to breeding sites, and are key to species recognition, mate choice, and the reinforcement of reproductive isolation (2,3,5–8). Through vocal signals, individuals convey information about their location, body size, and identity, allowing for effective social and reproductive interactions (2,3,5,9–12). Given the centrality of acoustic communication in frog biology, it is especially surprising that many species appear to thrive despite lacking the capacity to detect vocalizations.

Because acoustic signals are so pivotal in frog communication, their detection depends on a suite of specialized auditory structures, including the tympanic membrane, tympanic annulus, middle ear bones, and inner ear (13–16). Remarkably, despite their importance in detecting vocal signals, these auditory components have been independently lost at least 38 times across the frog phylogeny (17). These losses are phylogenetically widespread but non-random: they are disproportionately found in lineages that are ground-dwelling, diurnal, and miniaturized, with many species also exhibiting restricted geographic distributions and occupying high-elevation habitats (17,18). Despite these associations, the mechanisms driving the reduction of auditory structures remain poorly understood. One hypothesis suggests that these losses may reflect selection favouring alternative, yet unidentified traits or behaviours that compensate for the diminished capacity for acoustic communication (17). Such compensatory traits are presumed to be especially critical in mating contexts, where effective signalling remains essential (15,17). A plausible explanation is that the reduction of one sensory modality may be accompanied by enhancements in another, a sensory trade-off that could enable species to maintain functional signalling despite the absence of conventional auditory mechanisms (19–22).

Several cases of sensory trade-offs have been documented among different lineages. One well-characterized example is the bat *Rhinolophus hipposideros*, which possesses relatively small eyes and altered spectral sensitivity, consistent with adaptations to specialized echolocation (20). Similarly, boas and pythons exhibit a simple duplex retina composed of single cones and rods, in contrast to the more complex retinal structure of caenophidian snakes (23). This visual simplicity, however, is compensated by the evolution of a specialized infrared (IR) detection system: thermoreceptors located in the labial pits allow the snake to detect thermal radiation emitted by warm objects, facilitating prey capture, predator avoidance, and thermoregulation (21). Another example is found in subterranean mammals, such as mole rats. These animals have reduced visual systems and instead rely heavily on other sensory modalities, including tactile input, olfaction, and bone-conducted vibration, which are essential for navigation and foraging in dark, subterranean environments (24). In these examples, and others, enhancement of one modality compensates for a diminished reliance on vision. Our study proposes an inverse scenario: enhancement of the visual system may represent an adaptive response to the reduction or loss of auditory input.

The visual system is a crucial and extensively studied sensory modality, notable for its capacity to deliver extensive, real-time information essential for animal perception and behaviour. Eye size and shape directly influence optical properties, affecting both the quantity and quality of visual input (23,25). Because improvements in sensitivity, acuity, and temporal resolution often involve trade-offs, increasing eye size is generally required to enhance overall visual performance without sacrificing specific functions (25,26). Previous studies in vertebrates have demonstrated that the evolution of eye size is strongly influenced by ecological factors (22,26–34), a trend also observed in frogs (25). This suggests that frogs adapted to distinct ecological niches face unique optical constraints linked to variations in eye size. For instance, arboreal frogs tend to invest more in their eyes, exhibiting larger absolute and relative eye sizes, which are proposed to accommodate rapid visual processing during jumps and to achieve high visual precision in complex arboreal settings (25). By contrast, fossorial species exhibit minimal investment in eye size, presumably reflecting adaptations to dark or abrasive subterranean environments (25). Other ecological factors, such as activity period and reproductive strategies, also play a role. Nocturnal species often develop larger eyes to improve their sensitivity to dim light, while those breeding in vegetation or near lotic water may benefit from enhanced vision to overcome auditory challenges caused by the noise of rushing water (25,35).

Beyond anatomical investment, visual cues are critical in mediating both interspecific and intraspecific interactions, serving as essential signals across a wide range of ecological and social contexts (35), and may carry increased functional importance in species where other sensory modalities are reduced. Many frogs use conspicuous displays, such as posture changes, limb movements, or exposure of coloured body surfaces, to deter predators, attract prey, or communicate with conspecifics (35–39). These behaviours are often context-dependent and particularly prominent during courtship and agonistic encounters, with their form and frequency often modulated by whether the target is a potential mate or a rival (35). In species with limited auditory capacities, visual signals may compensate for the loss of acoustic communication, serving as a complementary channel for mate attraction and social interaction. This behavioural reliance on vision could impose selective pressures favouring increased investment in the visual system, particularly in lineages with auditory reductions. Building on this premise, we assess variation in visual system investment among frogs exposed to different ecological pressures, incorporating for the first time data on the condition of the auditory system. We hypothesize that the visual system compensates, at least in part, for the reduction of auditory perception. Accordingly, we predict that frog species with reduced or absent auditory structures will exhibit greater investment in their visual system, including the evolution of larger eyes to enhance visual capability and maintain effective communication and environmental perception in the absence of “conventional” hearing.

## Methodology

### Sampling and Morphometric Measurements

Starting from the morphometric dataset of Thomas et al. (25), which included 640 adult frog specimens from 220 frog species, we examined an additional 468 adult frog specimens from the amphibian collection at the QCAZ Museum of Zoology, Pontificia Universidad Católica del Ecuador. Specimens were assigned to 55 species across nine families: Aromobatidae, Bufonidae, Centrolenidae, Ceratophryidae, Craugastoridae, Hemiphractidae, Hylidae, Microhylidae, and Strabomantidae (see Tables S1 and S2). Our sampling emphasized species with reductions of the auditory system, which are particularly well represented in the QCAZ collection. We also included species with fully developed auditory structures to provide an appropriate baseline for comparison among datasets. In addition, we increased the number of individuals for some species that were represented by only a single specimen in the original dataset of Thomas et al. (25), thereby reducing sampling error and strengthening phylogenetic coverage.

Adulthood was determined by gonadal inspection: males were classified as adult if they had well-developed testes, and females if they had developed oocytes and/or convoluted oviducts (40). For each specimen, we recorded body mass (RM), snout–vent length (SVL), eye diameter (ED), and cornea diameter (CD) using a digital calliper (±0.01 mm) and a precision scale (±0.01 g), following Thomas et al. (25). Mean ED and CD were calculated per individual prior to analysis. These morphometric data allowed us to evaluate visual system investment in the context of auditory reduction and ecological variation. Eleven species overlapped between the two datasets and were counted only once, yielding a final dataset of 264 species that spans all 55 frog families and ensures broad ecological and phylogenetic representation (Table S3).

**Figure 1.**
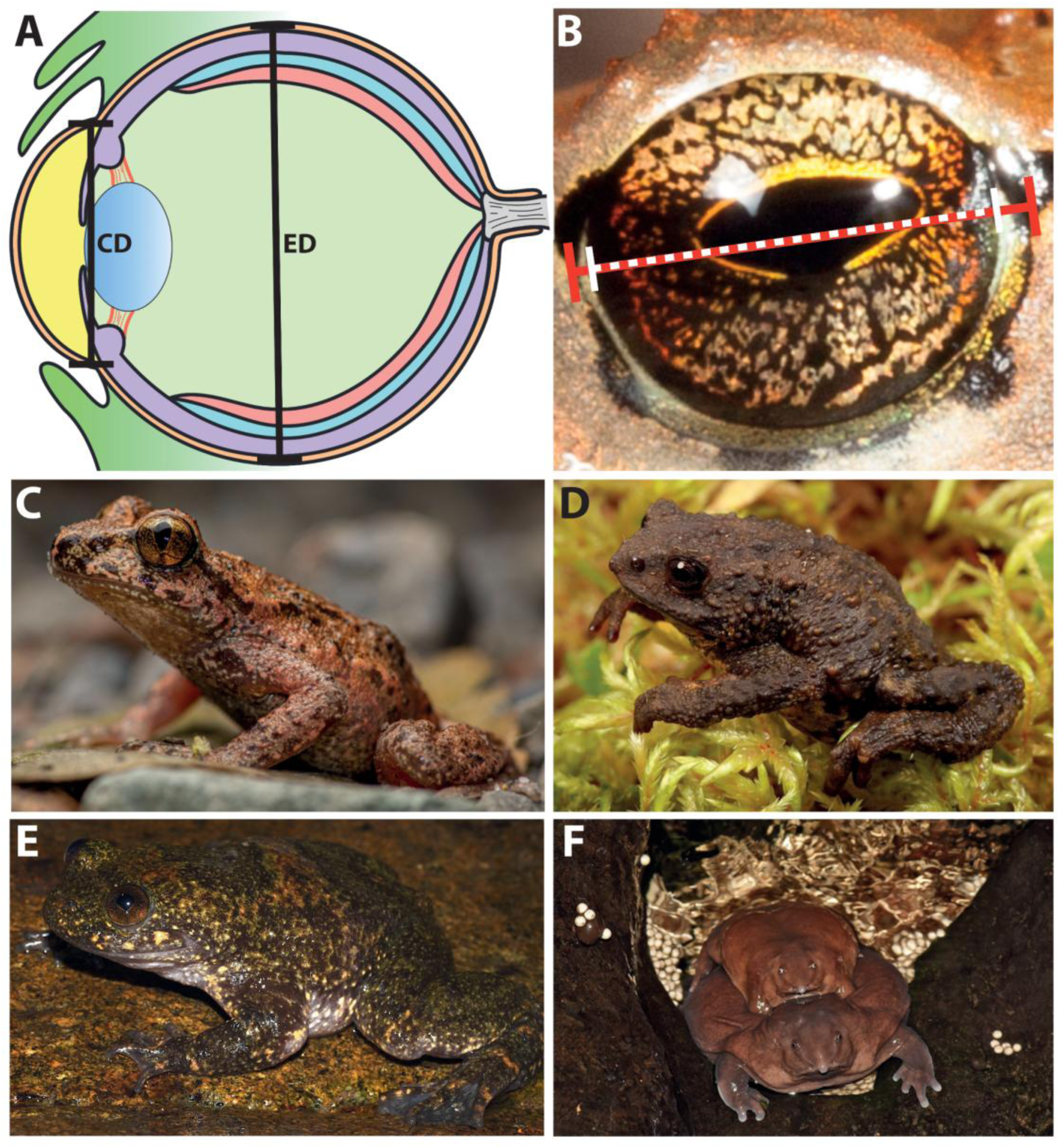
Examples of morphometric measurements and frogs with varying sensory system states. **(*A*)** Diagram of a frog eye illustrating the measured parameters: CD = Cornea Diameter; ED = Eye Diameter. Diagram by M. Fujita. **(*B*)** Eye measurements: red line indicates eye diameter; dashed white line, corneal diameter. **(*C*)** *Ascaphus truei*, semiaquatic species with reduction of the auditory system. Photo by Emma Steigerwald. **(*D*)** *Osornophryne guacamayo*, ground-dwelling species with reduction of the auditory system. Photo by S. R. Ron. **(*E*)** *Barbourula busuangensis*, acuatic species with reduction of the auditory system. Photo by: Ignacio de la Riva. **(*F*)** *Nasikabatrachus sahyadrensis*, fossorial species with reduction of the auditory system. Photo by: Sandeep Das.

### Phylogeny and Ecological Classification

To account for phylogenetic signal in our analysis, we aligned species with the most recent and comprehensive molecular phylogeny for amphibians (41), pruning it to include only the 264 species in our dataset. For 13 species not present in the tree, we substituted their closest congeneric relatives using the *ape* package in R (42). These replacements were incorporated as polytomies and subsequently resolved using the *chronos* function in *ape* (43–45). To maintain phylogenetic consistency, all substitutions and polytomies were restricted to species within the same genus.

We classified species according to five ecological traits: (1) status of the auditory system for hearing airborne sounds (maintained or lost); and five traits from Thomas et al. (25), (2) adult habitat (semiaquatic, aquatic, scansorial, ground-dwelling, fossorial, subfossorial), (3) activity period (diurnal, nocturnal, or both), (4) mating habitat (lotic water, lentic water, plants, or ground), and (5) life-history strategy (presence or absence of free-living larvae). Trait data were gathered from Thomas et al. (25), AmphibiaWeb, Bioweb, and primary literature, including species descriptions (see Table S1). Auditory system status was determined based on Womack and Hoke (17), who defined “earless” taxa as those lacking the columella, tympanic membrane, tympanic annulus, and middle ear cavity. The activity period classification was also updated following Womack and Hoke (17).

### Allometric patterns and ecological correlates in frog eye size

We first tested whether previously described allometric patterns (25) held when expanding the dataset to include more species with auditory reductions. We then examined whether auditory system loss correlated with increased investment in the visual system, and how this relationship might vary across ecological contexts.

We used phylogenetic generalized least squares (PGLS) models from the *caper* package (v. 1.0.3) (46) in R to assess allometric relationships between log-transformed species means of ED, CD, RM, and SVL. Specifically, we fit maximum-likelihood estimations of lambda (λ) to log-transformed species means for eye diameter (ED) *vs* snout-vent length (SVL), ED *vs* relative mass (RM), corneal diameter (CD) *vs* ED, CD *vs* SVL, CD *vs* RM, and SVL *vs* RM. To further evaluate allometric patterns, we performed ordinary least-squares (OLS) regressions using the *stats* package (v. 4.4.2) (47) and standardized major axis (SMA) regressions using the *smatr* package (v. 3.4-8) (48) in R, all based on the same log-transformed dataset.

To test differences in visual investment across ecological states, we used Wilcoxon rank sum tests or, for traits with more than two categories, Kruskal–Wallis tests using the *stats* package (v. 4.4.2) (47). We also implemented phylogenetic ANCOVAs via PGLS models using *caper* (v. 1.0.1) to examine the effect of each ecological trait on ED and CD while controlling for body size (RM or SVL). Separate models were run for each ecological variable to avoid overparameterization. Finally, two-way ANCOVAs via PGLS were used to evaluate the combined effects of two ecological traits on eye and cornea investments. Visualizations of eye size, corneal size, and ecological states on the phylogeny were generated using *ape* (v. 5.8) (42)and *phytools* (49). A full script to reproduce the analyses is available on Github (*link tbd*).

## Results

### Species with auditory reductions exhibit reduced absolute and relative eye size

Our analyses reveal that frogs with auditory reductions have consistently smaller eyes compared to species without auditory reduction, contrary to our initial prediction. This pattern holds true for both absolute eye diameter and eye size relative to body mass (eye investment). These differences remain significant even after controlling for confounding factors such as habitat (Fig. 3, Figs. S2–S4). Mean absolute eye diameter was 4.02 mm in species with auditory loss, compared to 5.98 mm in those without (Wilcoxon test: W = 3040.5, p < 0.001). Relative eye size, scaled by body mass and phylogenetically corrected, was also reduced in species with auditory loss (Wilcoxon test: W = 3144, p < 0.001), averaging 0.89× the expected value versus 1.07× in other species. Phylogenetic ANCOVA confirmed significant effects of both body mass (F_1,257_ = 1047.108, p < 0.001) and auditory loss (F_1,257_ = 20.365, p < 0.001) on eye size.

**Figure 2.**
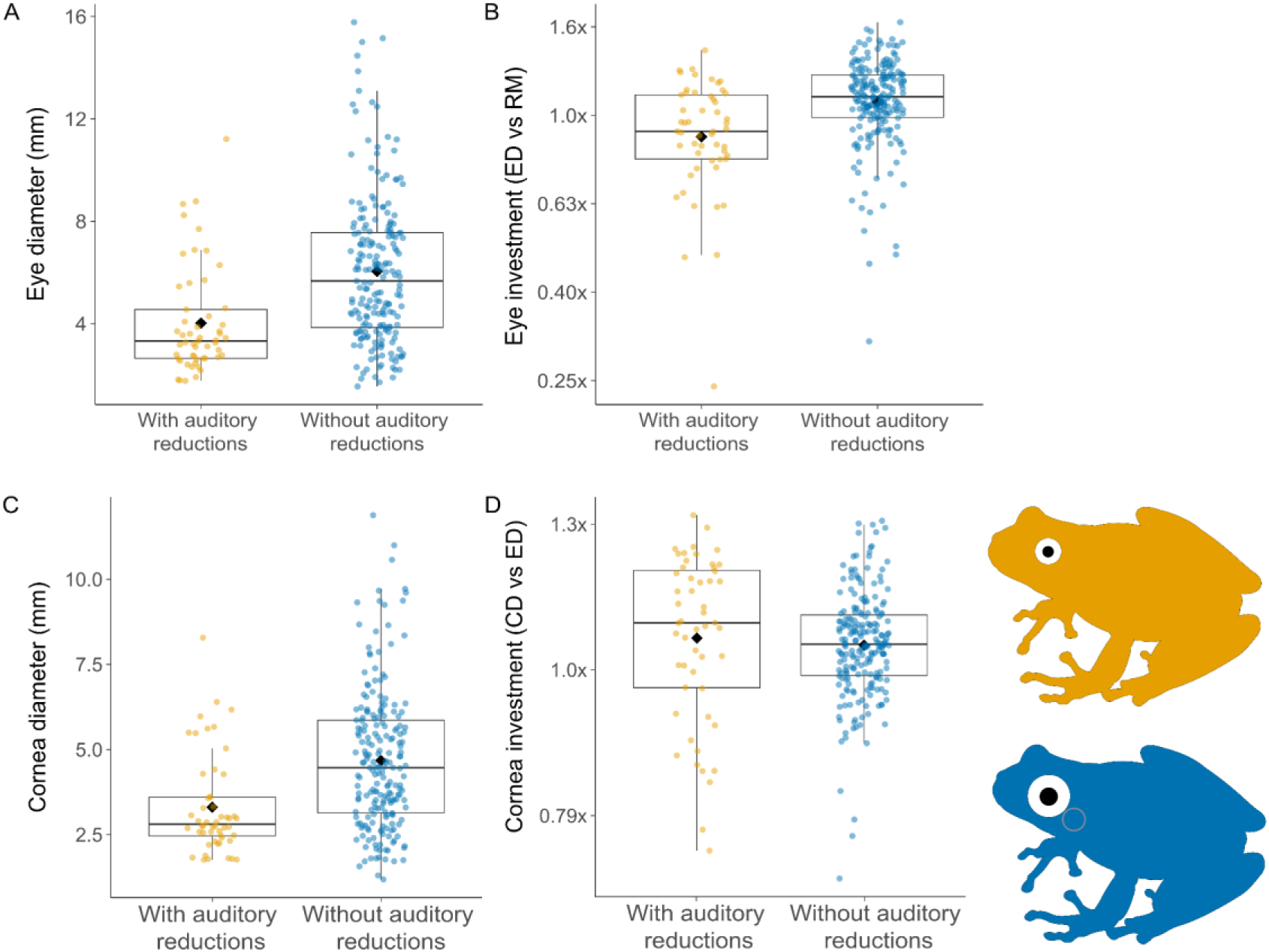
Visual investment across frogs with auditory reductions. **(*A*)** Absolute eye diameters and **(*B*)** eye investment across groups. **(*C*)** Absolute cornea diameter and (***D*)** relative corneal investment. All panels include data from all species in the dataset.

**Figure 3.**
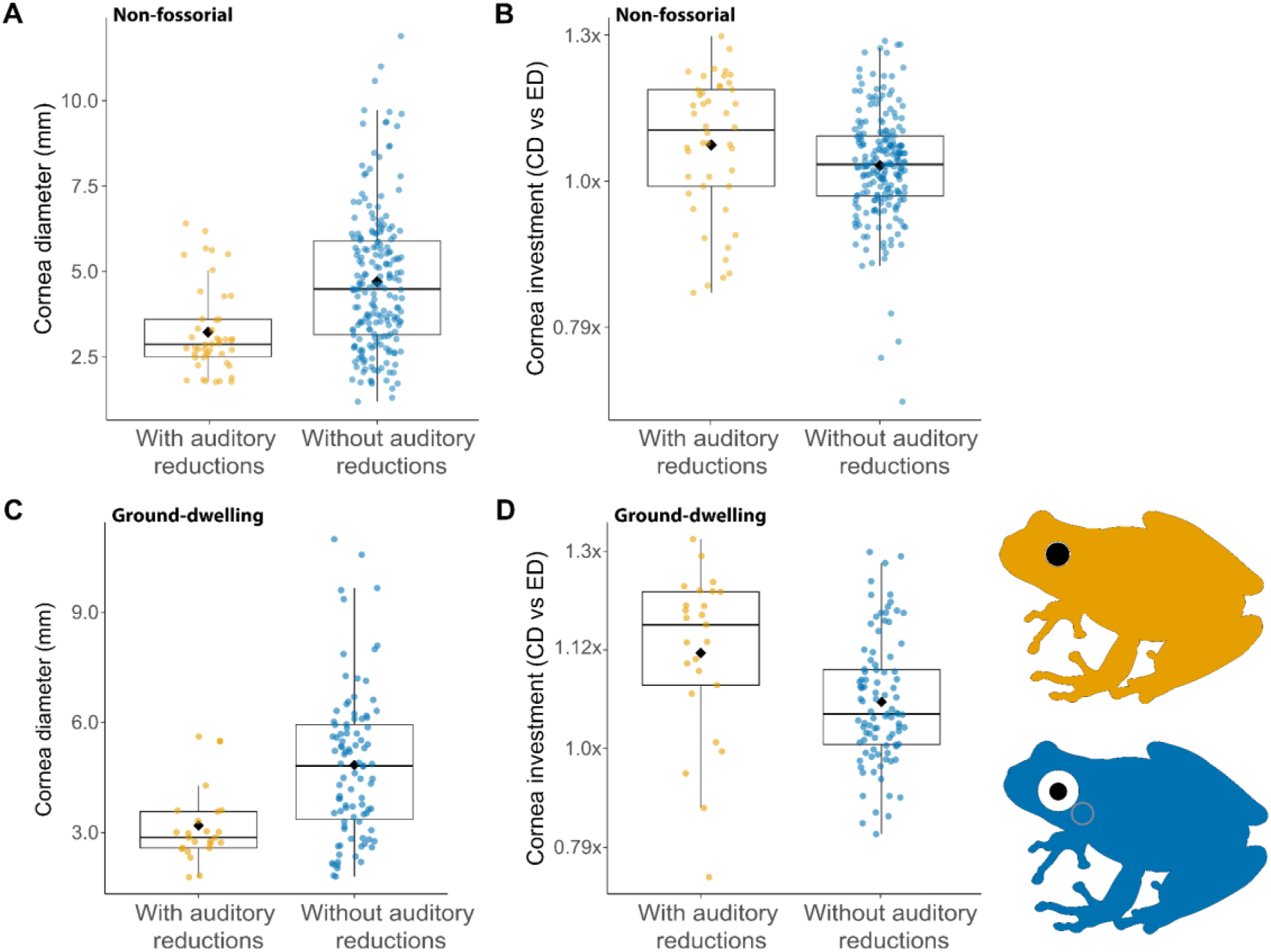
Cornea size and investment vary significantly across frogs with auditory reductions. **(A)** Absolute cornea diameter and **(B)** relative corneal investment, based on analyses excluding fossorial species (n = 254 spp.). (C) Absolute cornea diameter and (D) relative corneal investment, based on analyses restricted to ground-dwelling species (n = 120 spp.). Boxplots illustrate trait differences across sensory system conditions.

Because habitat strongly influences eye morphology, with fossorial and aquatic species typically exhibiting reduced eye size and investment (25), we tested whether these ecological factors could explain the observed pattern. A two-way phylogenetic ANCOVA revealed significant main effects of auditory system reduction (F_1,248_ = 20.462, p < 0.001) and adult habitat (F_1,248_ = 3.421, p = 0.005) on absolute eye size, but not significant interaction between them (F_1,248_ = 0.42, p = 0.833). Likewise, for relative eye size, both auditory loss (F_1,236_ = 27.428, p < 0.001) and adult habitat (F_5,236_ = 8.939, p < 0.001) had significant effects, with body mass exerting the strongest influence overall (F_1,236_ = 1307.736, p < 0.001). In addition, interactions between body mass and auditory reduction (F_1,236_ = 6.891, p = 0.009), as well as between auditory reduction and adult habitat (F_5,236_ = 3.412, p = 0.005) were also significant.

### Species with auditory loss have greater corneal investment

Given that species with auditory loss exhibit smaller eyes and reduced eye investment, we next examined corneal size and corneal investment. Across analyses, species with auditory loss consistently exhibited smaller absolute cornea diameters. By contrast, relative cornea investment tended to be higher in these species, though this difference was specific to non-fossorial taxa (Figs. 3–4, S2, S5).

**Figure 4.**
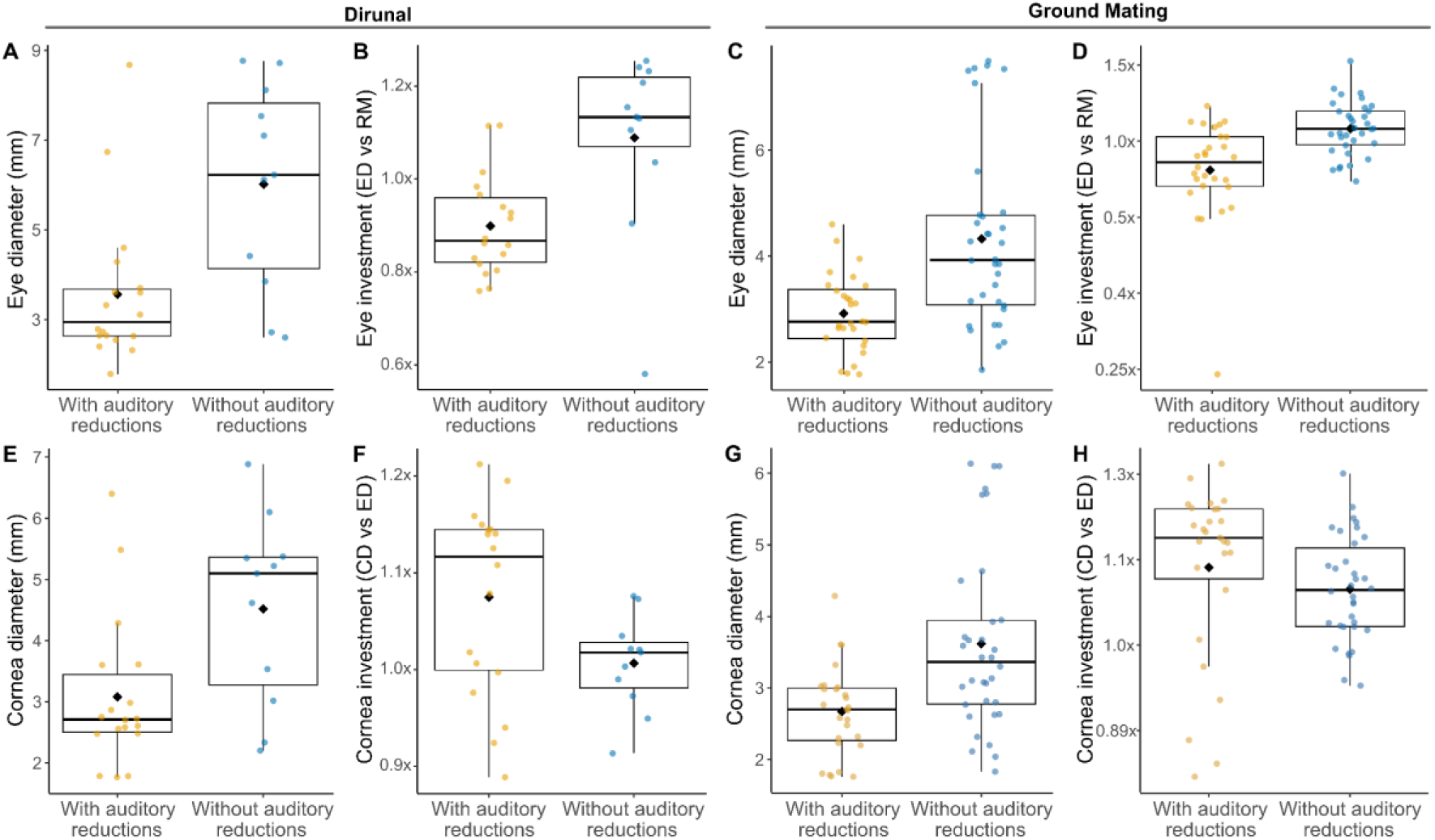
Eye and cornea size and investment in diurnal and ground-mating frog species. **(*A*)** Absolute eye diameter and **(*B*)** relative eye investment in diurnal species (n = 29 spp.). **(*C*)** Absolute eye diameter and **(*D*)** relative eye investment in species that mate on the ground (*n* = 63 spp.). **(*E*)** Absolute cornea diameter and **(*F*)** relative cornea investment in diurnal species (n = 29 spp.). **(*G*)** Absolute cornea diameter and **(*H*)** relative cornea investment in ground-mating species (*n* = 63 spp.). Boxplots show trait differences between species with and without auditory system reduction.

Similar to the pattern observed in eye size, species with auditory loss exhibited significantly smaller absolute cornea diameters (mean = 3.31 mm) compared to species without auditory reduction (mean = 4.68 mm; Wilcoxon test: W = 3061.5, p < 0.001). Relative cornea investment (*i.e.*, CD *vs* ED) showed no significant difference between species with auditory loss (mean = 1.05×) and those without (mean = 1.04×) (Wilcoxon test: W = 6286, p = 0.09). Phylogenetic ANCOVA revealed a significant main effect of eye size on cornea size (F_1,259_ = 4813.373, p < 0.001), but no significant main effect of auditory loss (F_1,259_ = 1.459, p = 0.228). A two-way phylogenetic ANCOVA testing the joint effects of adult habitat and sensory loss revealed significant main effects of both auditory loss (F_1,250_ = 23.477, p < 0.001) and habitat (F_5,250_ = 3.960, p = 0.002) on absolute cornea size, but not significant interaction between them (F_5,250_ = 0.698, p = 0.625). Similarly, for relative cornea investment, both auditory loss (F_1,238_ = 653.758, p < 0.001) and habitat (F_5,238_ = 130.628, p < 0.001) had significant effects, with eye size exerting the strongest overall influence (F_1,238_ = 5048.208, p < 0.001). In addition, the interaction between auditory reduction and adult habitat (F_5,238_ = 2.899, p = 0.015) was also significant.

Because the effect of auditory loss was only revealed when accounting for habitat variation, we suspected that not all species with auditory reduction exhibited the same pattern. In particular, fossorial species may be expected to be under different selective pressures and indeed several species with auditory loss are also fossorial. To further investigate this, we first excluded fossorial species and found that species with auditory loss still had significantly smaller absolute cornea diameters (mean = 3.84 mm) compared to those without auditory reduction (mean = 5.99 mm; Wilcoxon test: W = 2567, p < 0.001). By contrast, relative cornea investment was slightly, but significantly, greater in species with auditory loss (mean = 1.06×) compared to those without (mean = 1.02×; Wilcoxon test: W = 5907, p = 0.011). Phylogenetic ANCOVA revealed significant main effects of both eye size (F_1,249_ = 4707.502, p < 0.001) and auditory loss (F_1,249_ = 497.580, p < 0.001) on cornea size. Patterns for eye size remained the same (see SI).

To further assess the effects of auditory reduction on eye and corneal scaling while controlling for the influence of habitat, we restricted the analysis to ground-dwelling species only (n = 120), which was the habitat category with the highest proportion of species with auditory reductions (25 out of 53). Ground-dwelling species with auditory loss still exhibited significantly smaller eyes in both absolute (Wilcoxon test: W = 499.5, p < 0.001; mean = 3.63 mm vs. 6.17 mm) and relative terms (Wilcoxon test: W = 638, p = 0.0006; 0.97× vs. 1.1×). Phylogenetic ANCOVA again showed significant effects of body mass (F_1,113_ = 973.875, p < 0.001) and auditory loss (F_1,113_ = 167.637, p < 0.001). We also found that ground-dwelling species had significantly smaller absolute cornea diameters (mean = 3.19 mm) compared to species without auditory reduction (mean = 4.84 mm; Wilcoxon test: W = 559.5, p < 0.001). However, species with auditory loss showed significantly greater corneal investment (mean = 1.12×) than those without loss (mean = 1.05×; Wilcoxon test: W = 1700, p = 0.0009). Phylogenetic ANCOVA supported significant main effects of both eye size (F_1,117_ = 2855.795, p < 0.001) and auditory loss (F_1,117_ = 372.209, p < 0.001) on cornea size.

Because diurnality is overrepresented among species with sensory system reductions (16), we next tested the effects of activity period and auditory loss on eye morphology. A two-way phylogenetic ANCOVA revealed significant main effects of auditory system reduction (F_1,218_ = 13.146, p < 0.001) and activity period (F_2,218_ = 3.566, p = 0.030) on absolute eye size, but no significant interaction between them. Likewise, for relative eye size, both auditory loss (F_1,210_ = 80.399, p < 0.001) and activity period (F_2,210_ = 7.012, p = 0.001) had significant effects, with body mass exerting the strongest influence overall (F_1,210_ = 772.107, p < 0.001). In addition, the interaction between body mass and auditory reduction (F_1,210_ = 9.178, p = 0.003) was also significant. Similarly, a two-way phylogenetic ANCOVA revealed that both activity period (F_2,216_ = 5.169, p = 0.006) and auditory system reduction (F_1,216_ = 13.605, p < 0.001) significantly affected absolute cornea size, with no significant interaction between them. For relative cornea size, significant main effects were found for body mass (F_1,210_ = 3359.669, p < 0.001), auditory loss (F_1,210_ = 272.233, p < 0.001), and activity period (F_2,210_ = 83.356, p < 0.001), but no significant interactions were detected among these factors.

When restricting the analysis to diurnal species only (n = 29 spp., Fig. 4), we found that those with auditory reductions exhibited significantly smaller eyes in both absolute (Wilcoxon test: W = 39, p = 0.006; mean = 3.56 mm vs. 6.01 mm) and relative terms (Wilcoxon test: W = 30, p = 0.001; 0.89× vs. 1.25×). Phylogenetic ANCOVA confirmed that both auditory system reduction (F_1,25_ = 77.751, p < 0.001) and body mass (F_1,25_ = 149.374, p < 0.001) had significant effects on eye size. When analysing the cornea, we found that diurnal species with auditory reductions exhibited significantly smaller corneas in absolute size (Wilcoxon test: W = 51, p = 0.03; mean = 3.08 mm vs. 4.52 mm). However, differences in relative cornea size (1.09× in species with auditory loss vs. 1.00× in those without) were not statistically significant (Wilcoxon test: W = 141, p = 0.06). Phylogenetic ANCOVA confirmed that both auditory system reduction (F_1,26_ = 104.717, p < 0.001) and body mass (F_1,26_ = 347.899, p < 0.001) had significant effects on cornea size.

Given that mating habitat was previously shown to shape independently eye and cornea morphology (25) (see Supplementary Material), we next evaluated whether this ecological variable could also help account for the variation associated with sensory system reductions. To this end, we performed two-way phylogenetic ANCOVAs, testing the combined effects of auditory loss and mating habitat, along with post hoc Wilcoxon comparisons. Analyses revealed that absolute eye size was significantly influenced by both auditory system reduction (F_1,210_ = 16.469, p < 0.001) and mating habitat (F_3,210_ = 9.767, p < 0.001), with no evidence for an interaction between them. A similar pattern emerged for relative eye size: both auditory loss (F_1,202_ = 118.118, p < 0.001) and mating habitat (F_3,202_ = 26.244, p = 0.001) showed significant effects, while body mass had the strongest overall influence (F_1,202_ = 896.041, p < 0.001). Additionally, we detected significant interactions between body mass and auditory reduction (F_1,202_ = 16.237, p < 0.001), as well as a three-way interaction between auditory loss, mating habitat, and body mass (F_3,202_ = 3.701, p = 0.013) on eye size. For corneal morphology, both mating habitat (F_3,210_ = 6.496, p < 0.001) and auditory system reduction (F_1,210_ = 20.083, p < 0.001) had significant effects on absolute cornea size, while their interaction was not significant. For relative cornea size, we found significant main effects of mating habitat (F_3,202_ = 163.608, p < 0.001), auditory loss (F_1,202_ = 323.137, p < 0.001), and body mass (F_1,202_ = 3308.245, p < 0.001). Furthermore, both the interaction between mating habitat and auditory loss (F_3,202_ = 4.403, p = 0.005) and the three-way interaction among mating habitat, auditory loss, and body mass (F_3,202_ = 3.093, p = 0.028) were also significant.

Because our data show that most species with sensory system reductions mate on the ground, we repeated the analyses restricted to ground-mating species (n = 63 spp., Fig. 4). Within this subset, those with auditory reductions exhibited significantly smaller eyes in both absolute (Wilcoxon test: W = 223, p = 0.0003; mean = 2.92 mm vs. 4.33 mm) and relative terms (Wilcoxon test: W = 198, p < 0.001; 0.84× vs. 1.08×). Phylogenetic ANCOVA confirmed significant effects of auditory system reduction (F_1,58_ = 10.673, p < 0.001), body mass (F_1,58_ = 162.569, p < 0.001), and their interaction (F_1,58_ = 13.042, p = 0.001) on eye size. When analysing the cornea, we found that ground-mating species with auditory reductions had significantly smaller absolute cornea diameters (Wilcoxon test: W = 227.5, p = 0.0007; mean = 2.67 mm vs. 3.62 mm). Differences in relative cornea size were also significant (Wilcoxon test: W = 611, p = 0.03), with greater corneal investment observed in species with auditory loss (mean = 1.11×) compared to those without (mean = 1.07×). Phylogenetic ANCOVA again revealed significant effects of both auditory system reduction (F_1,58_ = 58.594, p < 0.001) and body mass (F_1,58_ = 1170.984, p < 0.001) on cornea size.

## Discussion

Our results show that frog species with auditory system reduction have consistently smaller eyes compared to those without auditory loss, both in absolute diameter and relative to body size. This pattern remained consistent across analyses, including when accounting for variation in habitat type and after excluding fossorial species that are expected to have smaller eyes. We also observed significant effects of both auditory loss and habitat on eye size, prompting additional analyses focused on ground-dwelling species, which were overrepresented among species with auditory reductions. The same pattern held: ground-dwelling species with auditory loss exhibited significantly smaller eyes. Species with auditory loss also had smaller absolute cornea diameters, but relative cornea investment (*i.e.*, cornea size relative to eye size) was significantly greater when fossorial species were excluded and when analyses were restricted to ground-dwelling taxa. These findings suggest that while overall eye size is constrained in species with auditory loss, specific eye components such as the cornea may be differentially affected under certain ecological contexts.

### Why prioritize investment in the cornea rather than the entire eye?

Our results highlight that reductions in the auditory system are closely linked to ecological context, particularly habitat, in shaping evolutionary investment in eye morphology among frogs. Many species with auditory loss inhabit fossorial or terrestrial environments, where ecological constraints strongly influence the utility and maintenance of visual structures. In fossorial species, the subterranean environment imposes strong limitations on vision, resulting in rudimentary visual systems characterized by consistently smaller eyes and corneas, a feature closely linked to their predominantly fossorial lifestyle (23,25,50,51). In such habitats, where visual cues are scarce, the metabolic and spatial costs of large eyes likely outweigh their benefits. Physical limitations such as darkness, abrasive substrates, and limited cranial space likely drive the evolutionary reduction of the visual system. This pattern is not exclusive to frogs but is also observed across other vertebrates, including mammals (52,53), fishes (54), reptiles (55), and caecilians (50). In these species, reliance likely shifts toward alternative sensory modalities such as chemoreception (56), mechanoreception (53), and, in some cases, bone-conducted auditory signals (28) rather than visual or airborne acoustic cues. It is likely due to these distinct evolutionary forces acting on fossorial species that we saw different scaling patterns in fossorial species with auditory reductions.

Our findings also indicate that most frogs with auditory system reductions are ground-dwelling, a pattern similarly reported by Womack and Hoke (17). These species are frequently found in leaf litter, among low vegetation, or near lotic water bodies, microhabitats that are structurally complex and visually cluttered. Although larger eyes can enhance visual acuity and sensitivity (25,57,58), they are metabolically costly to produce and maintain (59,60), and can pose biomechanical challenges, particularly for small-bodied, hopping animals. Consistent with this, our results, and those of Womack and Hoke (17), show that species with sensory reductions tend to be smaller in body size than those without, which may further constrain eye size due to allometric scaling. Moreover, large, protruding eyes increase cranial mass (61), raise the risk of mechanical injury from surrounding vegetation, and can hinder movement in cluttered environments (57), likely selecting against increased investment in eye structures under these ecological contexts.

Besides adult habitat and auditory loss, other ecological factors appear to influence eye size and related structures. Our analyses show that mating habitat and activity period significantly affect eye and cornea size alongside sensory loss. Frogs that mate primarily on the ground, overrepresented among species with auditory loss in our dataset, possess the smallest eyes in both absolute and relative terms, consistent with previous findings (25). Importantly, our results also reveal that these ground-mating species display increased relative corneal investment, suggesting a reallocation of visual resources despite an overall reduction in eye size. Regarding activity period, diurnal species, also overrepresented among auditory-loss frogs (17), generally invest less in eye size compared to nocturnal or mixed-activity species, a trend consistent with broader vertebrate patterns linking increased visual investment to the need for enhanced sensitivity in low-light environments (30,62). Although no significant differences in corneal size were detected among activity periods, our data reveal a main effect of activity period on both eye and cornea dimensions.

Given these ecological and anatomical constraints, reduced eye size associated with adult habitat, reproductive habitat, and diurnal activity, we propose that an increase in relative corneal size may be the optimal adaptive response for frogs with auditory losses. This potential adaptation could compensate for sensory reductions and their already small eyes, enhancing visual capabilities while minimizing metabolic and locomotive costs, thus offering a plausible solution to the environmental challenges these species face.

### Does having a larger cornea enhance the visual system?

Visual capacity is shaped by a variety of anatomical and optical factors beyond eye size. Structural differences, such as a higher rod-to-cone ratio (63), tubular-shaped eyes (57), and a larger cornea can significantly affect species-specific visual abilities (29,58). Although research on corneal size in frogs is limited, it is well-established in vertebrates more broadly that both the size and curvature of the cornea play crucial roles in determining visual sensitivity (23,58,64). The cornea’s size and curvature directly limit the maximum pupillary aperture, thereby controlling the amount of light entering the eye (58). While a larger cornea facilitates a greater pupillary aperture, enhancing retinal image brightness, this increased aperture size also broadens the field of vision, contributing to improved overall visual performance (23). To capture light rays reaching the anterior retina and extend the eye’s periscopic range, the cornea must be wider than the pupil (23). Additionally, a larger cornea and greater corneal investment increase retinal image brightness, enhancing visual sensitivity by maximizing the amount of light that reaches the retina (30,65). When other factors, such as eye size and corneal transmission, are held constant, an eye with a cornea occupying a larger proportion of the anterior surface can admit more light, thus improving visual sensitivity compared to an eye with a smaller corneal proportion (65). The optical properties of the cornea-lens system are optimized to maximize image brightness within a specific light range, independent of the overall size of the eye. The cornea-to-lens ratio governs the refractive power of the cornea, with a larger cornea typically corresponding to a proportionally larger lens (23,65). Studies conducted in nocturnal mammals and birds have shown that a larger relative cornea size is often coupled with a larger lens and a posterior nodal point shift toward the retina. In these species, this configuration of the dioptric system enables the production of a brighter retinal image by efficiently utilizing available light, thereby further enhancing visual sensitivity (58,65).

Beyond lens enlargement, corneal expansion may be accompanied by structural modifications and neural adaptations that enhance visual sensitivity, although these remain unexamined in species with auditory reductions. Retinal specializations, such as visual streaks or areas of elevated cell density along the naso-temporal axis, have been observed in both the ganglion cell layer (GCL) in *Boana raniceps*, *Bombina orientalis*, *Heleioporus eyrei* and *Rhinella marina* (66–70) and the inner nuclear layer (INL) in *Rhinella marina* and *Xenopus laevis* (71), suggesting adaptations for enhanced visual acuity (23,67). In *Rhinella marina* these streaks are also associated with a non-uniform distribution of photoreceptors, with peak densities along the naso-temporal axis (67). The spatial overlap between photoreceptor-rich regions and areas of high neuronal density in the GCL and INL defines a zone of acute vision, similar to the area centralis or visual streak found in other vertebrates (67). A comparable organization may exist in frogs with sensory reductions, potentially enhancing visual resolution in specific parts of the visual field. Complementing these structural changes, retinal modifications, such as the slenderness and spacing of visual cells, could further enhance both resolution and sensitivity (23). Narrower rods and cones might improve fine-detail detection, while closely spaced cells could reduce blurring of adjacent points of light. Elongation of rod outer segments may lower the stimulation threshold, allowing the retina to respond to dimmer light and potentially increasing overall visual sensitivity (23).

Finally, in frogs with reduced reliance on other sensory modalities, enhanced visual capacity may be accompanied not only by peripheral adaptations such as corneal enlargement and retinal specialization, but also by increased neural investment in brain regions dedicated to visual processing. In particular, these species may exhibit greater development of the optic tectum, the primary centre for visual integration in amphibians (72). This association between peripheral and central adaptations has also been documented in other vertebrate groups, suggesting a broader pattern of coordinated investment across the visual system (73,74), but remains unstudied in frogs.

## Conclusion

Frogs predominantly use acoustic communication for a wide range of social interactions, with mate attraction and selection being among the most critical functions. In species that have lost the ability to detect airborne sounds, this essential communication channel is impaired, necessitating compensatory adaptations. Contrary to our initial expectation that these species would evolve larger eyes to improve visual signalling, we found that they instead exhibit a relative enlargement of the cornea alongside an overall reduction in eye size. This corneal expansion likely enhances visual sensitivity, allowing these frogs to better utilize visual cues for social and reproductive behaviours while avoiding the metabolic and locomotor costs of larger eyes. Such an adaptation is particularly important for small-bodied species facing anatomical constraints. Our results highlight the role of sensory compensation following auditory loss and provide new insights into the evolutionary dynamics of communication strategies in frogs.

## Notes

### Competing Interest Statement

The authors have declared no competing interest.

### Funding

- Natural Sciences and Engineering Research Council: Dicovery Grant (to RKS)

## References

1. Wilkins MR, Seddon N, Safran RJ. Evolutionary divergence in acoustic signals: causes and consequences. Trends Ecol Evol. 2013 Mar;28(3):156–66.

2. Gerhardt HC, Huber F, Simmons AM. Acoustic Communication in Insects and Anurans: Common Problems and Diverse Solutions. J Acoust Soc Am. 2003 Aug 1;114(2):559–559.

3. Wells KD. The Ecology and Behavior of Amphibians [Internet]. University of Chicago Press; 2007 [cited 2024 Mar 12]. Available from: http://www.bibliovault.org/BV.landing.epl?ISBN=9780226893341

4. Köhler J, Jansen M, Rodríguez A, Kok PJR, Toledo LF, Emmrich M, et al. The use of bioacoustics in anuran taxonomy: theory, terminology, methods and recommendations for best practice. Zootaxa [Internet]. 2017 Apr 11 [cited 2024 Dec 12];4251(1). Available from: https://www.mapress.com/zt/article/view/zootaxa.4251.1.1

5. Toledo LF, Martins IA, Bruschi DP, Passos MA, Alexandre C, Haddad CFB. The anuran calling repertoire in the light of social context. Acta Ethologica. 2015 Jun;18(2):87–99.

6. Endler JA. Signals, Signal Conditions, and the Direction of Evolution. Am Nat. 1992 Mar;139:S125–53.

7. Kelley DB. Vocal communication in frogs. Curr Opin Neurobiol. 2004 Dec;14(6):751–7.

8. Wells KD, Schwartz JJ. The Behavioral Ecology of Anuran Communication. In: Narins PM, Feng AS, Fay RR, Popper AN, editors. Hearing and Sound Communication in Amphibians [Internet]. Springer New York; 2006 [cited 2024 Dec 12]. p. 44–86. (Springer Handbook of Auditory Research; vol. 28). Available from: http://link.springer.com/10.1007/978-0-387-47796-1_3

9. Bee MA, Gerhardt HC. Neighbour–stranger discrimination by territorial male bullfrogs (Rana catesbeiana): I. Acoustic basis. Anim Behav. 2001 Dec;62(6):1129– 40.

10. Davis MS. Acoustically mediated neighbor recognition in the North American bullfrog, Rana catesbeiana. Behav Ecol Sociobiol. 1987 Sep;21(3):185–90.

11. Ryan M. ANURAN COMMUNICATION. Copeia. 2002 Feb;2001(1):252–4.

12. Simmons AM. Call recognition in the bullfrog, *Rana catesbeiana* : Generalization along the duration continuum. J Acoust Soc Am. 2004 Mar 1;115(3):1345–55.

13. Barry TH. The ontogenesis of the sound-conducting apparatus of Bufo angusticeps Smith. Morphol Jahrb. 1956;97:477–544.

14. Helff OM. Studies on Amphibian Metamorphosis. III. The Influence of the Annular Tympanic Cartilage on the Formation of the Tympanic Membrane. Physiol Zool. 1928 Oct;1(4):463–95.

15. Pereyra MO, Womack MC, Barrionuevo JS, Blotto BL, Baldo D, Targino M, et al. The complex evolutionary history of the tympanic middle ear in frogs and toads (Anura). Sci Rep. 2016 Sep 28;6(1):34130.

16. Sedra SN, Michael MI. The ontogenesis of the sound conducting apparatus of the egyptian toad, bufo regularis reuss, with a review of this apparatus in salientia. J Morphol. 1959 Mar;104(2):359–75.

17. Womack MC, Hoke KL. Convergent anuran middle ear loss lacks a universal, adaptive explanation. Brain Behav Evol [Internet]. 2023 Nov 1 [cited 2023 Nov 16]; Available from: https://karger.com/doi/10.1159/000534936

18. Von May R, Lehr E, Rabosky DL. Evolutionary radiation of earless frogs in the Andes: molecular phylogenetics and habitat shifts in high-elevation terrestrial breeding frogs. PeerJ. 2018 Feb 22;6:e4313.

19. Mason MJ, Narins PM. Vibrometric studies of the middle ear of the bullfrog *Rana catesbeiana* I. The extrastapes. J Exp Biol. 2002 Oct 15;205(20):3153–65.

20. Gutierrez EDA, Schott RK, Preston MW, Loureiro LO, Lim BK, Chang BSW. The role of ecological factors in shaping bat cone opsin evolution. Proc R Soc B Biol Sci. 2018 Apr 11;285(1876):20172835.

21. Emer SA, Grace MS, Mora CV, Harvey MT. Pit organ-based infrared discrimination sensitivity and signal transduction in the Burmese python (Python molurus bivitattus). Behav Brain Res. 2022 Jul;429:113910.

22. Martínez-Martínez CA, Martins HOJ, Kobal ROAC, Cordeiro GD, Hrncir M, Alves-dos-Santos I. Unique morphological and morphometric traits of nocturnal bee antennae. Apidologie. 2024 Dec;55(6):78.

23. Walls GL. The vertebrate eye and its adaptive radiation [by] Gordon Lynn Walls. [Internet]. Bloomfield Hills, Mich.,: Cranbrook Institute of Science,; 1942 [cited 2024 Dec 12]. Available from: http://www.biodiversitylibrary.org/bibliography/7369

24. Mason MJ, Narins PM. Seismic sensitivity in the desert golden mole (Eremitalpa granti): A review. J Comp Psychol. 2002;116(2):158–63.

25. Thomas KN, Gower DJ, Bell RC, Fujita MK, Schott RK, Streicher JW. Eye size and investment in frogs and toads correlate with adult habitat, activity pattern and breeding ecology. Proc R Soc B Biol Sci. 2020 Sep 30;287(1935):20201393.

26. Land MF, Nilsson DE. Animal eyes. 2nd ed. Oxford: Oxford university press; 2012. (Oxford animal biology series).

27. Brooke MDL, Hanley S, Laughlin SB. The scaling of eye size with body mass in birds. Proc R Soc Lond B Biol Sci. 1999 Feb 22;266(1417):405–12.

28. Castiglione GM, Chiu YLI, Gutierrez EDA, Van Nynatten A, Hauser FE, Preston M, et al. Convergent evolution of dim light vision in owls and deep-diving whales. Curr Biol. 2023 Nov;33(21):4733–4740.e4.

29. Hall MI. Comparative analysis of the size and shape of the lizard eye. Zoology. 2008 Jan;111(1):62–75.

30. Kirk E. Effects of activity pattern on eye size and orbital aperture size in primates. J Hum Evol. 2006 Aug;51(2):159–70.

31. Liu Y, Ding L, Lei J, Zhao E, Tang Y. Eye size variation reflects habitat and daily activity patterns in colubrid snakes. J Morphol. 2012 Aug;273(8):883–93.

32. Mitkus M, Potier S, Martin GR, Duriez O, Kelber A. Raptor Vision. In: Oxford Research Encyclopedia of Neuroscience [Internet]. Oxford University Press; 2018 [cited 2024 Dec 12]. Available from: http://oxfordre.com/neuroscience/view/10.1093/acrefore/9780190264086.001.0001/acrefore-9780190264086-e-232

33. Schmitz L, Wainwright PC. Nocturnality constrains morphological and functional diversity in the eyes of reef fishes. BMC Evol Biol. 2011;11(1):338.

34. Werner YL, Seifan T. Eye size in geckos: Asymmetry, allometry, sexual dimorphism, and behavioral correlates. J Morphol. 2006 Dec;267(12):1486–500.

35. Hödl W, Amézquita A. Visual signaling in anuran amphibians. In: Anuran communication. Ryan, M.J. Washington, DC: Smithsonian Inst. Press.; 2001. p. 121–41.

36. Bajger J. Diversity of defensive responses in populations of Fire Toads (Bombina bombina and Bombina variegata). Herpetologica. 1980;36:133–7.

37. Edmunds M. Visual Signaling in Anuran Amphibians. In: Anuran communication. Ryan MJ. Washington, USA: Smithsonian lust. Press; 2001. p. 121–41.

38. Hinsche G. Vergleichende untersuchungen zum sogenannten Unkenreflex. Biol Zentralblatt. 1926;46:296–305.

39. Myers C. The distribution and behavior of a tropical horned frog, Cerathyla panamensis Stejneger. Herpetologica. 1966;22:68–71.

40. Duellman WE, Lehr E. Terrestrial-breeding frogs (Strabomantidae) in Peru. Munster, Germany: Nature und Tier Verlag.; 2009.

41. Portik DM, Streicher JW, Wiens JJ. Frog phylogeny: A time-calibrated, species-level tree based on hundreds of loci and 5,242 species. Mol Phylogenet Evol. 2023 Nov;188:107907.

42. Paradis E, Claude J, Strimmer K. APE: Analyses of Phylogenetics and Evolution in R language. Bioinformatics. 2004 Jan 22;20(2):289–90.

43. Kim J, Sanderson MJ. Penalized Likelihood Phylogenetic Inference: Bridging the Parsimony-Likelihood Gap. Syst Biol. 2008 Oct 1;57(5):665–74.

44. Paradis E. Molecular dating of phylogenies by likelihood methods: A comparison of models and a new information criterion. Mol Phylogenet Evol. 2013 May;67(2):436–44.

45. Sanderson MJ. Estimating Absolute Rates of Molecular Evolution and Divergence Times: A Penalized Likelihood Approach. Mol Biol Evol. 2002 Jan 1;19(1):101–9.

46. Orme D, Freckleton R, Thomas G, Petzoldt T, Fritz S, Isaac N, et al. caper: Comparative Analyses of Phylogenetics and Evolution in R [Internet]. 2011 [cited 2025 Jun 6]. p. 1.0.3. Available from: https://CRAN.R-project.org/package=caper

47. R Core Team. R: a language and environment for statistical computing. Vienna, Austria: R Foundation for Statistical Computing. 2019;

48. Warton DI, Duursma RA, Falster DS, Taskinen S. smatr 3– an R package for estimation and inference about allometric lines. Methods Ecol Evol. 2012 Apr;3(2):257–9.

49. Revell LJ. phytools: an R package for phylogenetic comparative biology (and other things). Methods Ecol Evol. 2012 Apr;3(2):217–23.

50. Mohun SM, Davies WL, Bowmaker JK, Pisani D, Himstedt W, Gower DJ, et al. Identification and characterization of visual pigments in caecilians (Amphibia: Gymnophiona), an order of limbless vertebrates with rudimentary eyes. J Exp Biol. 2010 Oct 15;213(20):3586–92.

51. Mohun SM, Wilkinson M. The eye of the caecilian *Rhinatrema bivittatum* (Amphibia: Gymnophiona: Rhinatrematidae). Acta Zool. 2015 Apr;96(2):147–53.

52. Borghi CE, Giannoni SM, Roig VG. Eye reduction in subterranean mammals and eye protective behavior in Ctenomys. Mastozool Neotrop. 2002;9:123–34.

53. Catania K. Mole senses. Curr Biol. 2009;29(17):R825–8.

54. Eagderi S, Adriaens D. Cephalic morphology of *Pythonichthys macrurus* (Heterenchelyidae: Anguilliformes): specializations for head-first burrowing. J Morphol. 2010 Sep;271(9):1053–65.

55. Yovanovich CAM, Pierotti MER, Rodrigues MT, Grant T. A dune with a view: the eyes of a neotropical fossorial lizard. Front Zool. 2019 Dec;16(1):17.

56. Martín J, Raya García E, Ortega J, López P. How to maintain underground social relationships? Chemosensory sex, partner and self recognition in a fossorial amphisbaenian. Bartos L, editor. PLOS ONE. 2020 Aug 19;15(8):e0237188.

57. Thomas RJ, Székely T, Powell RF, Cuthill IC. Eye size, foraging methods and the timing of foraging in shorebirds. Funct Ecol. 2006 Feb;20(1):157–65.

58. Lisney TJ, Iwaniuk AN, Kolominsky J, Bandet MV, Corfield JR, Wylie DR. Interspecifc variation in eye shape and retinal topography in seven species of galliform bird (Aves: Galliformes: Phasianidae). J Comp Physiol A. 2012 Oct;198(10):717–31.

59. Laughlin SB, De Ruyter Van Steveninck RR, Anderson JC. The metabolic cost of neural information. Nat Neurosci. 1998 May;1(1):36–41.

60. Moran D, Softley R, Warrant EJ. The energetic cost of vision and the evolution of eyeless Mexican cavefish. Sci Adv. 2015 Sep 4;1(8):e1500363.

61. Laughlin SB. Towards the cost of seeing. Nervous Systems and Behaviour. Vis Res. 1995;36:1529–41.

62. Garamszegi LZ, Møller AP, Erritzøe J. Coevolving avian eye size and brain size in relation to prey capture and nocturnality. Proc R Soc Lond B Biol Sci. 2002 May 7;269(1494):961–7.

63. Waldvogel JA. The Bird’s Eye View. Am Sci. 1990 Agosto;78(4):342–53.

64. Martin GR. Schematic Eye Models in Vertebrates. In: Ottoson D, Autrum H, Perl ER, Schmidt RF, Shimazu H, Willis WD, editors. Progress in Sensory Physiology [Internet]. Berlin, Heidelberg: Springer Berlin Heidelberg; 1983 [cited 2024 Dec 12]. p. 43–81. (Progress in Sensory Physiology; vol. 4). Available from: http://link.springer.com/10.1007/978-3-642-69163-8_2

65. Kirk EC. Comparative morphology of the eye in primates. Anat Rec A Discov Mol Cell Evol Biol. 2004 Nov;281A(1):1095–103.

66. Bousfield JD, Pessoa VF. Changes in ganglion cell density during post-metamorphic development in a neotropical tree frog Hyla raniceps. Vision Res. 1980 Jan;20(6):501–10.

67. Zhang Y, Straznicky C. The morphology and distribution of photoreceptors in the retina of Bufo marinus. Anat Embryol (Berl) [Internet]. 1991 Jan [cited 2025 Apr 19];183(1). Available from: http://link.springer.com/10.1007/BF00185840

68. Dunlop SA, Beazley LD. Changing retinal ganglion cell distribution in the frog *Heleioporus eyrei*. J Comp Neurol. 1981 Oct 20;202(2):221–36.

69. Nguyen VS, Straznicky C. The development and the topographic organization of the retinal ganglion cell layer in Bufo marinus. Exp Brain Res [Internet]. 1989 Apr [cited 2025 Apr 19];75(2). Available from: http://link.springer.com/10.1007/BF00247940

70. Pushchin II, Zyumchenko NE. Retinal ganglion cell topography and spatial resolving power in the oriental fire-bellied toad *Bombina orientalis*. J Integr Neurosci. 2015 Dec;14(04):521–34.

71. Zhu B, Hiscock J, Straznicky C. The changing distribution of neurons in the inner nuclear layer from metamorphosis to adult: a morphometric analysis of the anuran retina. Anat Embryol (Berl) [Internet]. 1990 Jul [cited 2025 Apr 19];181(6). Available from: http://link.springer.com/10.1007/BF00174630

72. Dong W, Lee RH, Xu H, Yang S, Pratt KG, Cao V, et al. Visual Avoidance in *Xenopus* Tadpoles Is Correlated With the Maturation of Visual Responses in the Optic Tectum. J Neurophysiol. 2009 Feb;101(2):803–15.

73. Yopak KE, Lisney TJ. Allometric Scaling of the Optic Tectum in Cartilaginous Fishes. Brain Behav Evol. 2012;80(2):108–26.

74. Barton RA. Visual specialization and brain evolution in primates. Proc R Soc Lond B Biol Sci. 1998 Oct 22;265(1409):1933–7.

