## Supplementary Information for "Eyes wide open: Frogs with reduction of auditory communication ability show lower eye, but higher corneal, investment"

#### Supporting Information:

Supplemental Results
Supplemental References
Tables S1-S3
Figures S1-S5

#### Supplemental Results

##### Scaling relationships: Anuran corneas scale hypoallometrically with eye and body size

For the analysis between anuran ED *vs.* RM, PGLS revealed a strong positive correlation, with a slope of 0.8532 ($p < 2.2e-16$, ±0.02735 SE) and a phylogenetic $\lambda$ of 0.958 (95% CI: 0.92–

0.98), indicating significant phylogenetic signal. The model explained 79.05% of the variance ($R^2 = 0.7905$, $p < 2.2e-16$). Similarly, OLS regression showed a slope of 0.8789 ($p < 2e-16$, (±0.0323 SE) and $R^2 = 0.741$ ($p < 2.2e-16$), further confirming the positive relationship between the traits. SMA analysis resulted in a slope of 1.0087 (95% CI: 0.9583–1.0619) and $R^2 = 0.741$ ($p < 2.22e-16$), indicating a near-isometric relationship between root mass and eye size, **with no significant deviation from isometry** ($p = 0.7379$).

Similarly, for the analysis between anuran ED *vs.* SVL, PGLS analysis revealed a significant positive correlation ($R^2 = 0.8204$, $p < 2.2e-16$), with a slope of 0.8554 (±0.0247 SE) and a strong phylogenetic signal ($\lambda = 0.953$; 95% CI: 0.913–0.976). OLS regression yielded a similar result, estimating a slope of 0.898 ($p < 2e-16$, ±0.0289 SE) with $R^2 = 0.7857$ ($p < 2.2e-16$). SMA analysis produced a slope of 0.9865 (95% CI: 0.941–1.0341) and $R^2 = 0.7857$ ($p < 2.22e-16$). A test of whether the SMA slope deviates from 1 yielded a p-value of 0.5719, indicating **no significant deviation from isometry**.

For the relationship between CD *vs.* ED, PGLS analysis revealed a significant positive correlation ($R^2 = 0.9491$, $p < 2.2e-16$), with a slope of 0.8859 ($p < 2.2e-16$, ± 0.0127 SE). The phylogenetic signal was moderate, with a $\lambda$ value of 0.674 (95% CI: 0.452–0.815, $p < 2.2e-16$). OLS regression produced a similar slope of 0.8871 ($p < 2.2e-16$, ± 0.0118 SE), with $R^2 = 0.9556$ ($p < 2.2e-16$), further supporting the observed correlation. SMA analysis estimated a slope of 0.9074 (95% CI: 0.8844–0.9311) and $R^2 = 0.9556$, ($p < 2.2e-16$) significantly lower than 1 ($p < 2.22e-16$) **indicating negative allometry in corneal investment relative to eye size**.

For the relationship between CD *vs*. RM, PGLS analysis revealed a significant positive correlation between cornea size and root mass ($R^2 = 0.7661$, $p < 2.2e-16$), with a slope of 0.7788 ($p < 2.2e-16$, ± 0.0269 SE). The phylogenetic signal was also strong with a $\lambda$ of 0.949 (95% CI: 0.903–0.974, $p < 2.2e-16$). OLS analysis produced similar results, with a slope of 0.7637 ($p < 2.2e-16$, ± 0.0326 SE), $R^2 = 0.6812$ ($p < 2.2e-16$), supporting the negative allometric relationship. SMA analysis estimated a slope of 0.8893 (95% CI: 0.8388–0.9428), with an $R^2$ of 0.6812 and significatively lower than 1 (p-value < 2.22e-16), further confirming the **negative allometric scaling between cornea size and root mass.**

Regarding the relationship between CD *vs.* SVL, the PGLS analysis revealed a significant positive correlation ($R^2 = 0.8204$, $p < 2.2e-16$), with a slope of 0.7894 ($p < 2.2e-16$, ± 0.0229 SE), indicating a negative allometric relationship between these variables. The $\lambda$ value was 0.958 (95% CI: 0.92–0.98, $p < 2.2e-16$), indicating a strong phylogenetic signal. The OLS analysis also showed a negative allometric relationship, with a slope of 0.7935 ($p < 2.2e-16$, ± 0.0285 SE), $R^2 = 0.7481$, and $p < 2.2e-16$. Similarly, the SMA analysis estimated a slope of 0.8671 (95% CI: 0.8260–0.9102), with an $R^2$ of 0.7481 and a p-value < 2.22e-16, confirming the **negative allometric relationship between cornea size and SVL.**

Finally, in the relationship between SVL *vs.* RM, the PGLS analysis revealed a significant positive correlation ($R^2 = 0.935$, $p < 2.2e-16$), with a slope of 0.9894 ($p < 2.2e-16$, ± 0.0136 SE), consistent with an isometric scaling pattern. The $\lambda$ value was 0.879 (95% CI: 0.743–0.945, $p < 2.2e-16$), indicating a strong phylogenetic signal. OLS results showed a slope of

0.9845 ($p < 2.2e-16$, ± 0.0128 SE) and $R^2 = 0.9577$ ($p < 2.2e-16$). SMA analysis estimated a slope of 1.006 (95% CI: 0.9809–1.0317), $R^2 = 0.9577$,and not significantly different from 1 ($p = 0.6388$), indicating **no significant deviation from isometry.**

#### Species with auditory reductions exhibit reduced absolute and relative eye size

##### ***Analysis restricted to fossorial species***

Given the strong influence of habitat, and the fact that several species with auditory loss are also fossorial, we repeated the analyses excluding fossorial species to determine whether the effect persisted independently of ecological specialization. After removing fossorial taxa, the pattern remained consistent: species with auditory loss still had significantly smaller eyes (Fig. S3), with mean absolute eye diameter of 3.84 mm compared to 5.99 mm (Wilcoxon test: $W = 2423$, $p < 0.001$), and lower relative eye size (0.92× vs. 1.03×; Wilcoxon test: $W = 2916$, $p < 0.001$). Phylogenetic ANCOVA further supported significant effects of both body mass ($F_{1,247} = 1319.110$, $p < 0.001$) and auditory loss ($F_{1,257} = 8.664$, $p = 0.004$).

##### ***Analysis analyzing the effect of sensory reductions and life history strategy***

When considering life-history strategy, a two-way phylogenetic ANCOVA revealed that only auditory system reduction had a significant effect on absolute eye size ($F_{1,235} = 25.118$, $p < 0.001$), while neither life-history strategy ($F_{1,235} = 3.190$, $p = 0.075$) nor its interaction with auditory loss ($F_{1,235} = 0.706$, $p = 0.401$) were significant. For relative eye size, significant main effects were found for auditory system reduction ($F_{1,231} = 117.813$, $p < 0.001$), life-history strategy ($F_{1,231} = 8.788$, $p = 0.003$), and body mass ($F_{1,231} = 950.204$, $p < 0.001$). Additionally, significant two-way interactions were detected between auditory loss and life-history strategy ($F_{1,231} = 9.275$, $p = 0.003$), auditory loss and body mass ($F_{1,231} = 4.258$, $p = 0.040$), and life-history strategy and body mass ($F_{1,231} = 4.000$, $p = 0.047$), whereas the three-way interaction was not significant ($F_{1,231} = 0.002$, $p = 0.967$). When examining absolute cornea size, auditory system reduction had a significant main effect ($F_{1,235} = 27.159$, $p < 0.001$), while life-history strategy ($F_{1,235} = 2.014$, $p = 0.157$) and its interaction with auditory loss ($F_{1,235} = 0.481$, $p = 0.489$) were not significant. Regarding relative cornea investment, strong main effects were observed for life-history strategy ($F_{1,231} = 82.253$, $p < 0.001$), auditory system reduction ($F_{1,231} = 537.440$, $p < 0.001$), and eye size ($F_{1,231} = 3778.520$, $p < 0.001$). However, none of the interaction terms reached statistical significance, including those between auditory reduction and life-history strategy ($F_{1,231} = 0.006$, $p = 0.938$), life-history strategy and eye size ($F_{1,231} = 0.010$, $p = 0.922$), auditory reduction and eye size ($F_{1,231} = 1.545$, $p = 0.215$), or the three-way interaction ($F_{1,231} = 0.139$, $p = 0.709$).

Given the apparent overrepresentation of direct developers among species with sensory system reductions, we next assessed whether this life history strategy could help explain the observed variation. To do so, we restricted our analyses to species without a free-living larval stage and examined differences in eye and corneal morphology within this subset.

When restricting the analysis to species without a free-living larval stage, we found that those with auditory reductions exhibited significantly smaller eyes in absolute size (Wilcoxon test: W = 82, p = 0.0001; mean = 2.79 mm vs. 4.77 mm), but not in relative size (Wilcoxon test: W = 158, p = 0.05; 0.95× vs. 1.05×). Phylogenetic ANCOVA showed that both auditory system reduction ($F_{1,45} = 10.128$, $p = 0.003$) and body mass ($F_{1,45} = 189.947$, $p < 0.001$) had significant effects on eye size, while their interaction was not significant ($F_{1,45} = 0.153$, $p = 0.697$). When examining the cornea, species without a free-living larva and with auditory reductions exhibited significantly smaller absolute cornea sizes (Wilcoxon test: W = 95, p = 0.0006; mean = 2.55 mm vs. 3.95 mm). However, differences in relative cornea size (1.1× in species with sensory loss vs. 1.08× in those without) were not statistically significant (Wilcoxon test: W = 313, p = 0.14). Phylogenetic ANCOVA confirmed significant effects of both auditory system reduction ($F_{1,46} = 88.595$, $p < 0.001$) and body mass ($F_{1,46} = 1078.942$, $p < 0.001$) on cornea size.

#### Other ecological factors influence anuran eye and cornea size and investments independently of auditory reductions

##### *Adult Habitat influences Eye and Corneal Investment*

Eye absolute size (Kruskal–Wallis $\chi^2 = 34.626$, df = 5, $p < 0.0001$) and eye investment (Kruskal–Wallis $\chi^2 = 61.743$, df = 5, $p < 0.0001$) showed significant differences across habitat categories. Consistent with the findings of Thomas et al., semi-aquatic species exhibited the largest absolute eye size (mean = 8.14 mm), followed by ground-dwelling (5.6 mm), scansorial (5.18 mm), fossorial (5.1 mm), aquatic (4.9 mm), and subfossorial species, which had the smallest eyes (mean = 4.2 mm). Regarding eye investment, scansorial (1.16×) and semi-aquatic species (1.17×) exhibited the highest levels of eye investment, followed by ground-dwelling species (1.07×). The lowest values were observed in subfossorial (0.89×), aquatic (0.71×), and fossorial species (0.64×). The phylogenetic ANCOVA found that both body mass ($F_{1,248} = 1188.555$, $p < 0.0001$) and adult habitat ($F_{5,248} = 10.812$, $p < 0.0001$) had significant positive effects on eye size. Additionally, the interaction between mass and adult habitat ($F_{5,248} = 2.775$, $p = 0.019$) also significantly influenced eye size.

Adult habitat significantly influenced both absolute and relative cornea size (Kruskal–Wallis $\chi^2 = 37.78$, df = 5, $p < 0.0001$; $\chi^2 = 26.06$, df = 5, $p < 0.0001$). Semi-aquatic species had the largest corneas (mean = 6.13 mm), followed by ground-dwelling (4.50 mm), scansorial (4.22 mm), fossorial (3.88 mm), aquatic (3.58 mm), and subfossorial species (3.30 mm). For relative cornea investment, ground-dwelling species showed the highest values (1.07×), followed by scansorial (1.06×), semi-aquatic (1.03×), subfossorial (1.00×), aquatic (0.94×), and fossorial species (0.92×). Phylogenetic ANCOVA confirmed significant effects of eye size ($F_{1,250}$ =

5743.22, $p < 0.0001$), adult habitat ($F_{5,250} = 4.68$, $p = 0.001$), and their interaction ($F_{5,250} = 2.68$, $p = 0.022$) on absolute cornea size.

***Mating habitats affect Eye and Corneal Investment***

Eye absolute size (Kruskal–Wallis $\chi^2 = 60.755$, df = 3, p = < 0.0001) and eye investment (Kruskal–Wallis $\chi^2 = 35.983$, df = 3, p = < 0.0001) also showed significant differences across mating habitats categories. Species that mate on the ground have the smallest eyes (mean = 3.69 mm), followed by species that mate on plants (5.26 mm), lentic water species (6.31 mm), and those that mate in lotic water have the largest eyes (7.71 mm). A similar pattern is observed for eye investment, where species that mate on the ground show the lowest investment (0.96x), followed by those that mate in lentic water (0.98x), on plants (1.16x), and the highest investment is found in species that mate in lotic water (1.19x). In the phylogenetic ANCOVA, both root mass ($F_{1,210} = 847.820$, p < 0.0001) and mating habitat ($F_{3,210} = 5.841$, p = 0.001) had significant positive effects on eye size. However, the interaction between eye size and mating habitat was not significant ($F_{3,210} = 1.356$, p = 0.26).

Mating habitat also significantly affected corneal traits (absolute: $\chi^2 = 47.22$, df = 3, $p < 0.0001$; relative: $\chi^2 = 29.49$, df = 3, $p < 0.0001$). Species mating on the ground had the smallest corneas (mean = 3.19 mm), followed by those mating on plants (4.25 mm), in lentic water (4.77 mm), and in lotic water (5.95 mm). In contrast, ground-mating species exhibited the highest cornea investment (1.09×), followed by lotic (1.05×), plant (1.04×), and lentic (1.00×) species. Phylogenetic ANCOVA showed that both eye size ($F_{1,213} = 3870.02$, $p < 0.0001$) and mating habitat ($F_{3,213} = 3.89$, $p = 0.01$) significantly influenced cornea size.

***Activity Period influences Eye and Cornea Investment***

Absolute eye size differed slightly but significantly among activity period categories (Kruskal–Wallis $\chi^2 = 6.234$, df = 2, p = 0.04), with exclusively diurnal species having the smallest eyes (mean = 4.49 mm), followed by exclusively nocturnal species (mean = 5.46 mm), and species with mixed activity patterns (both diurnal and nocturnal) showing the largest eyes (mean = 6.04 mm). In contrast, relative eye investment did not differ significantly among activity categories (Kruskal–Wallis $\chi^2 = 5.800$, df = 2, p = 0.055). However, in both studies, the phylogenetic ANCOVA detected significant effects of body size ($F_{1,218} = 725.837$, p < 0.0001) and activity period ($F_{2,218} = 14.22$, p < 0.0001) on eye size.

Activity period did not significantly affect corneal traits (Kruskal–Wallis: absolute: $\chi^2 = 5.57$, df = 2, $p = 0.062$; relative: $\chi^2 = 1.13$, df = 2, $p = 0.57$). However, phylogenetic ANCOVA revealed significant effects of activity period ($F_{2,218} = 83.67$, $p < 0.0001$) and eye size ($F_{2,218} = 3635.08$, $p < 0.0001$) on absolute cornea size.

***Life History strategies affect Corneal Investment but not Eye Investment***

We found significant differences in absolute eye size (Wilcoxon test: W = 6839.5, $p < 0.0001$), but not in eye investment (Wilcoxon test: W = 5118, p = 0.2838), across life history strategies. Species with free-living larvae had larger absolute eye sizes (mean = 6.06 mm) than those without (mean = 4.21 mm). Although no differences were found in relative investment, the phylogenetic ANCOVA revealed a significant main effect only of body size ($F_{1,235} = 930.451$, $p < 0.0001$).

Life history strategy significantly influenced corneal dimensions. Species with free-living larvae had larger absolute corneas (mean = 4.73 mm) than those without (3.55 mm) (Wilcoxon test: W = 6521.5, $p < 0.0001$). However, cornea investment was higher in species without free-living larvae (1.10× vs. 1.03×) (Wilcoxon test: W = 2834, $p < 0.0001$). These patterns were supported by phylogenetic ANCOVA (eye size: $F_{1,236} = 5002.12$, $p < 0.0001$; life history: $F_{1,236} = 14.14$, $p < 0.0001$).

#### Supplemental References


AmphibiaWeb.org

Andersson, Lars Gabriel (1903) (https://www.biodiversitylibrary.org/part/28991)

Biju, S. D., S. Garg, K. V. Gururaja, Y. S. Shouche, and S. A. Walujkar. 2014. DNA barcoding reveals unprecedented diversity in Dancing Frogs of India (Micrixalidae, Micrixalus): a taxonomic revision with description of 14 new species. Ceylon Journal of Science. Biological Sciences 43: 1—87.

Bioweb.bio

Boulenger, G. A. 1882. Catalogue of the Batrachia Salientia s. Ecaudata in the Collection of the British Museum. Second Edition. London: Taylor and Francis.

Boulenger, G. A. 1897. Descriptions of new lizards and frogs from Mount Victoria, Owen Stanley Range, New Guinea, collected by Mr. A. S. Anthony. Annals and Magazine of Natural History, Series 6, 19: 6–13.

Boulenger, G. A. 1903. Descriptions of three new batrachians from Tonkin. Annals and Magazine of Natural History, Series 7, 12: 186–188.

Boulenger, G. A. 1905. Descriptions of new West-African frogs of the genera Petropedetes and Bulua. Annals and Magazine of Natural History, Series 7, 15: 281–283.

Boulenger, G. A. 1909. Descriptions of three new frogs discovered by Dr. P. Krefft in Usambara, German East Africa. Annals and Magazine of Natural History, Series 8, 4: 496–497.

Chan, K. O., M. A. Muin, M. S. S. Anuar, J. Andam, N. Razak, and M. A. Aziz. 2019. First checklist on the amphibians and reptiles of Mount Korbu, the second highest peak in Peninsular Malaysia. Check List. The Journal of Biodiversity Data 15: 1055–1069 (https://doi.org/10.15560/15.6.1055).

Chuaynkern, Y., et al. 2010. A revision of species in the subgenus Nidirana Dubois, 1992, with special attention to the identity of specimens allocated to Rana adenopleura Boulenger, 1909, and Rana chapaensis (Bourret, 1937) (Amphibia: Anura: Ranidae) from Thailand and Laos. Raffles Bulletin of Zoology 58: 291–310.

Coloma, L. A., and W. E. Duellman. 2025. Amphibians of Ecuador. Volume 2. Pipidae, Telmatobidae, Microhylidae, Dendrobatidae, Ranidae, Bufonidae, and Hylidae. Boca Raton, FL: CRC Press.

de Sá, R. O., T. Grant, A. Camargo, W. R. Heyer, M. L. Ponssa, and E. L. Stanley. 2014. Systematics of the Neotropical genus Leptodactylus Fitzinger, 1826 (Anura: Leptodactylidae): Phylogeny, the relevance of non-molecular evidence, and species accounts. South American Journal of Herpetology 9(Spec. Issue 1): 1–128.

Du Preez, L. H., and V. C. Carruthers. 2017. Frogs of Southern Africa: A Complete Guide. Cape Town, South Africa: Struik Nature.

Dubois, A. 1980. Notes sur la systematique et la repartition des amphibiens anoures de Chine et des regions avoisinantes IV. Classification generique et subgenerique des Pelobatidae Megophryinae. Bulletin Mensuel de la Société Linnéenne de Lyon 49: 469–482.

Duellman, W. E., & Lynch, J. D. (1969). Descriptions of Atelopus Tadpoles and Their Relevance to Atelopodid Classification. Herpetologica, 25(4), 231–240. http://www.jstor.org/stable/3891213

Duellman, W. E., and E. Lehr. (2009). Terrestrial-breeding frogs (Strabomantidae) in Peru. Münster, Germany: Nature und Tier Verlag.

Dunn, E. R. 1948. American frogs of the family Pipidae. American Museum Novitates 1384: 1–13 (http://digitallibrary.amnh.org/handle/2246/4379).

Elliot, L., H. C. Gerhardt, and C. Davidson. 2009. The Frogs and Toads of North America. A Comprehensive Guide to Their Identification, Behavior, and Calls. Boston and New York: Houghton Mifflin Harcourt.

Fouquet, A., K. Leblanc, A.-C. Fabre, M. T. Rodrigues, M. Menin, E. A. Courtois, M. Dewynter, M. Hölting, R. Ernst, P. L. V. Peloso, and P. J. R. Kok. 2021. Comparative osteology of the fossorial frogs of the genus Synapturanus (Anura, Microhylidae) with the description of three new species from the Eastern Guiana Shield. Zoologischer Anzeiger 293: 46–73 (https://doi.org/10.1016/j.jcz.2021.05.003).

Gillespie, G., E. Ahmad, and A. Shia. 2021. Field Guide to the Frogs of the Lower Kinabatangan Region, Sabah. Sabah, Malaysia: Hutan.

Günther, A. C. L. G. 1873. Description of two new species of frogs from Australia. Annals and Magazine of Natural History, Series 4, 11: 349–350.

Heyer, W. R. 1971. Mating calls of some frogs from Thailand. Fieldiana. Zoology 58: 61–82., Dubois, A., A. Ohler, and R. A. Pyron. 2021. New concepts and methods for phylogenetic taxonomy and nomenclature in zoology, exemplified by a new ranked cladonomy of recent amphibians (Lissamphibia). Megataxa 5: 1–738 (https://doi.org/10.11646/megataxa.5.1.1).

Heyer, W. R. 1971. Mating calls of some frogs from Thailand. Fieldiana. Zoology 58: 61–82.

Inger, R. F. 1966. The systematics and zoogeography of the Amphibia of Borneo. Fieldiana. Zoology 52: 1–402.

Inger, R. F. 1970. A new species of frog of the genus Rana from Thailand. Fieldiana. Zoology 51: 169–174.

Inger, R. F., and J. D. Romer. 1961. A new pelobatid frog of the genus Megophrys from Hong Kong. Fieldiana. Zoology 39: 533–538.

Kok, P. J. R., and M. Kalamandeen. 2008. Introduction to the Taxonomy of the Amphibians of Kaieteur National Park, Guyana. Abc Taxa: A Series of Manuals Dedicated to Capacity Building in Taxonomy and Collection Management, volume 5. Brussels, Belgium: Belgian Development Corporation.

Matsui, M. 1997. Call characteristics of Malaysian Leptolalax with a description of two new species (Anura: Pelobatidae). Copeia 1997: 158–165.

Meneses, C. G., C. D. Siler, P. A. Alviola, J. B. Balatibat, J. C. T. Gonzalez, C. A. Natividad, and R. M. Brown. 2022. Amphibian and reptile diversity along a ridge-to-reef elevational gradient on a small isolated oceanic island of the central Philippines. Check List. The Journal of Biodiversity Data 18: 941–984 (https://doi.org/10.15560/18.5.941).

Mkonyi (2020) (https://doi.org/10.1080/00222933.2020.1728410)

Mueses-Cisneros et al. (2012) (10.11646/zootaxa.3447.1.2)

Nunes-de-Almeida, C. H. L., C. L. de Assis, R. N. Feio, and L. F. Toledo. 2016. Redescription of the advertisement call of five species of Thoropa (Anura, Cycloramphidae), including recordings of rare and endangered species. PLoS (Public Library of Science) One 11(9: e0162617): 1–12 (doi:10.1371/journal.pone.0162617).

Ohler, A. 2003. Revision of the genus Ophryophryne Boulenger, 1903 (Megophryidae) with description of two new species. Alytes. Paris 21: 23–42.

Ortega-Andrade, H. M., Rojas-Soto, O. R., Espinosa de los Monteros, A., Valencia, J. H., Read, M., & Ron, S. R. (2017). Revalidation of Pristimantis brevicrus (Anura, Craugastoridae) with taxonomic comments on a widespread Amazonian direct-developing frog. Herpetological Journal, 31(2), 81–97.

Parker, H. W. 1934. A Monograph of the Frogs of the Family Microhylidae.  London: Trustees of the British Museum.

Peters, J. A. (1973). The frog genus Atelopus in Ecuador (Anura: Bufonidae). Smithsonian Contributions to Zoology 145:1-49.

Poyarkov, N. A., Jr., T. V. Duong, N. L. Orlov, S. I. Gogoleva, A. B. Vassilieva, L. T. Nguyen, V. D. H. Nguyen, S. N. Nguyen, J. Che, and S. Mahony. 2017. Molecular, morphological and acoustic assessment of the genus Ophryophryne (Anura, Megophryidae) from Langbian Plateau, southern Vietnam, with description of a new species. ZooKeys 672: 49–120.,

Smith, M. A. 1921. New or little-known reptiles and batrachians from southern Annam (Indo-China). Proceedings of the Zoological Society of London 1921: 423–440.

Springer, L. E., and C. M. Schalk. 2016. Lepidobatrachus laevis. Catalogue of American Amphibians and Reptiles 904: 1–16.

Stuart, S. N., Hoffmann, M., Chanson, J., Cox, N., Berridge, R., Ramani, P., Young, B. E. (2008). Threatened Amphibians of the World. Lynx Edicions. España 160.

Stuart, S. N., M. Hoffmann, J. Chanson, N. Cox, R. Berridge, P. Ramani, and B. Young eds., . 2008. Threatened Amphibians of the World. Barcelona, Spain; International Union for the Conservation of Nature, Gland. Switzerland; Conservation International, Arlington, Virginia, U.S.A.: Lynx Editions.

Stuart, S. N., M. Hoffmann, J. Chanson, N. Cox, R. Berridge, P. Ramani, and B. Young eds., . 2008. Threatened Amphibians of the World. Barcelona, Spain; International Union for the

Conservation of Nature, Gland. Switzerland; Conservation International, Arlington, Virginia, U.S.A.: Lynx Editions.

Sumarli, A. X. Y., L. L. Grismer, M. S. S. Anuar, M. A. Muin, and E. S. H. Quah. 2015. First report on the amphibians and reptiles of a remote mountain, Gunung Tebu in northeastern Peninsular Malaysia. Check List. The Journal of Biodiversity Data 11(4, Art. 1679): 1–16.

Thomas, K. N., Gower, D. J., Bell, R. C., Fujita, M. K., Schott, R. K., & Streicher, J. W. (2020). Eye size and investment in frogs and toads correlate with adult habitat, activity pattern and breeding ecology. Proceedings of the Royal Society B: Biological Sciences, 287(1935), 20201393. https://doi.org/10.1098/rspb.2020.1393

Tyler, M. J., M. M. Davies, and A. A. Martin. 1981. Australian frogs of the leptodactylid genus Uperoleia Gray. Australian Journal of Zoology, Supplemental Series 29 (79): 1–64.

Weygoldt, P. 1976. Beobachtungen zur Biologie und Ethologie von Pipa (Hemipipa) carvalhoi Mir. Rib. 1937. (Anura, Pipidae). Zeitschrift für Tierpsychologie 40: 80–99.

Wogan, G. O. U., K. S. Lwin, H. Win, and T. Thin. 2004. The advertisement call of Brachytarsophrys feae (Boulenger 1887) (Anura: Megophryidae). Proceedings of the California Academy of Sciences 55: 251–254.

Womack, M. C., & Hoke, K. L. (2023). Convergent anuran middle ear loss lacks a universal, adaptive explanation. Brain, Behavior and Evolution. https://doi.org/10.1159/000534936

#### Tables S1-S3

**Table S1. Specimens measured in this study and in Thomas et al. (2020).** (INDEPENDENT FILE)

**Table S2. Species included in this study, including those from Thomas et al. (2020).** The table presents the ecological traits analyzed and the character states for each species. (INDEPENDENT FILE)

**Table S3. Overview of sampling effort.** The table includes all extant anuran families sampled in this study. (INDEPENDENT FILE)

### Figures S1-S8

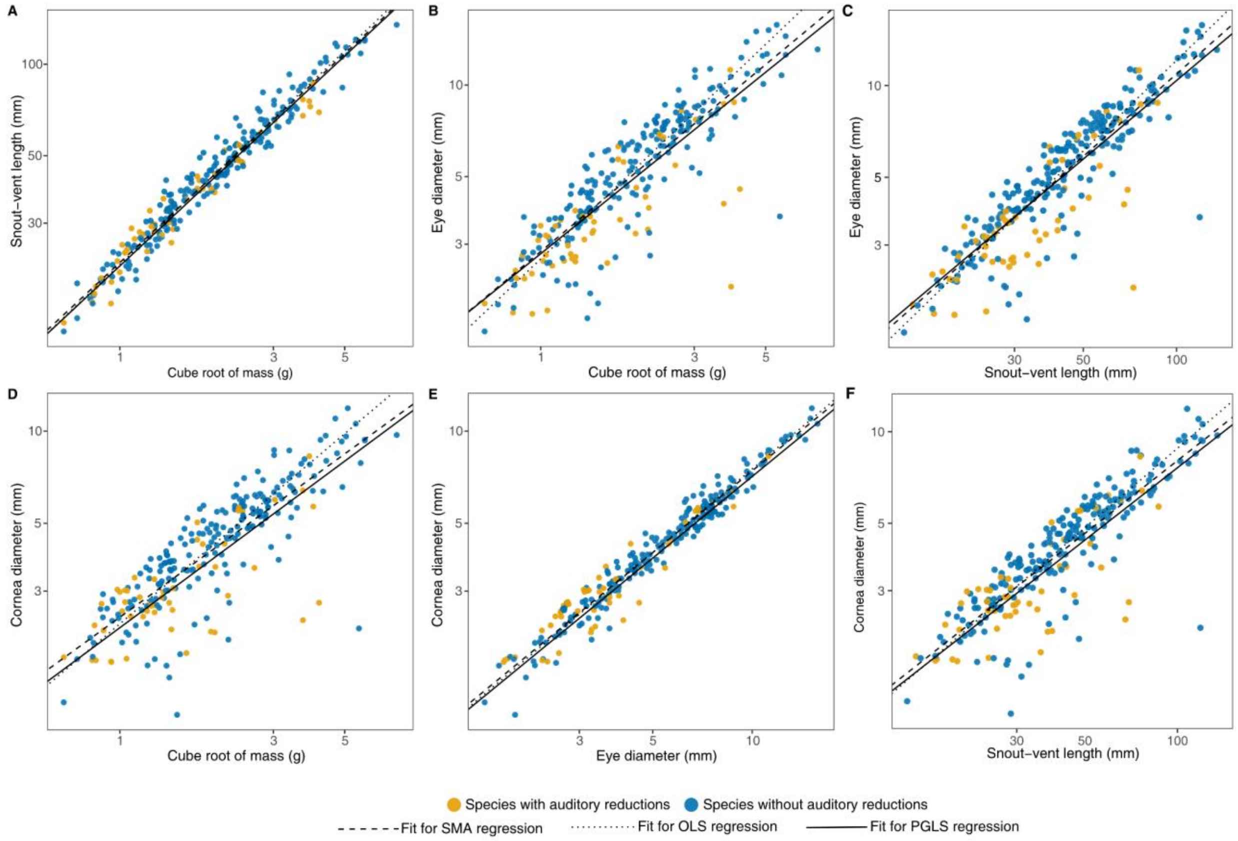


**Figure S1. Scaling of eyes and corneas of different frog species. (*A*)** Snout–vent length (SVL) scales isometrically with the cube root of body mass (RM), reflecting the high correlation between these two-size metrics. **(*B*)** Eye diameter (ED) scales isometrically with RM and **(*C*)** SVL. **(*D*)** Corneal diameter (CD) shows negative allometry with RM, **(*E*)** ED, **(*F*)** and SVL. Fitted lines for each analysis are shown. Species are color-coded based on the condition of their sensory systems.

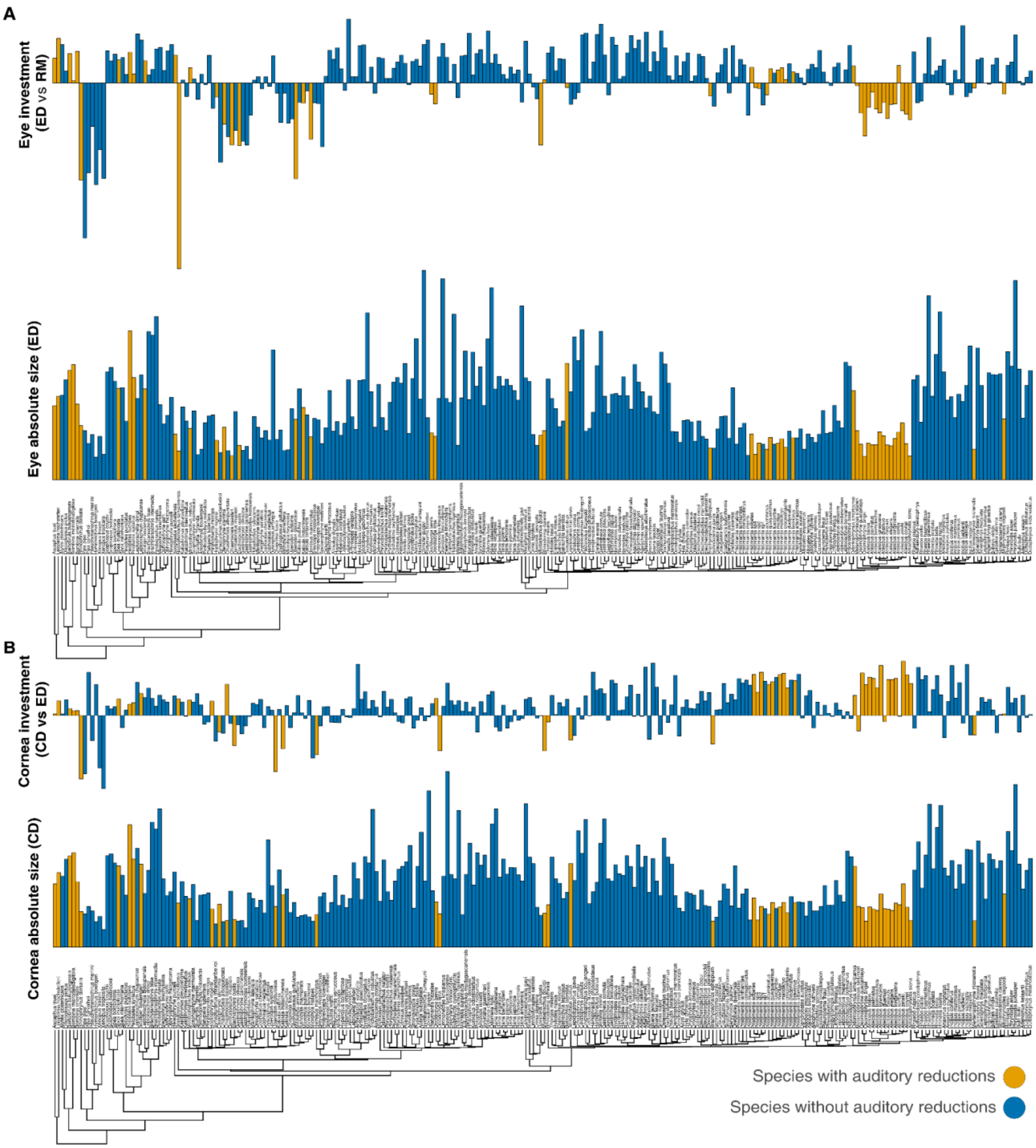


**Figure S2. Species with auditory loss exhibit reduced visual investment, having significantly smaller absolute and relative eye sizes than species without loss.** (***A***) Phylogeny, adapted from Portik et al. (2023), shows species means for absolute eye diameter (ED) and relative eye investment (ED *vs* RM) and (***B***) species mean values for absolute cornea diameter (CD) and relative corneal investment (CD vs ED). Analyses were performed using the full dataset, including all species regardless of habitat.

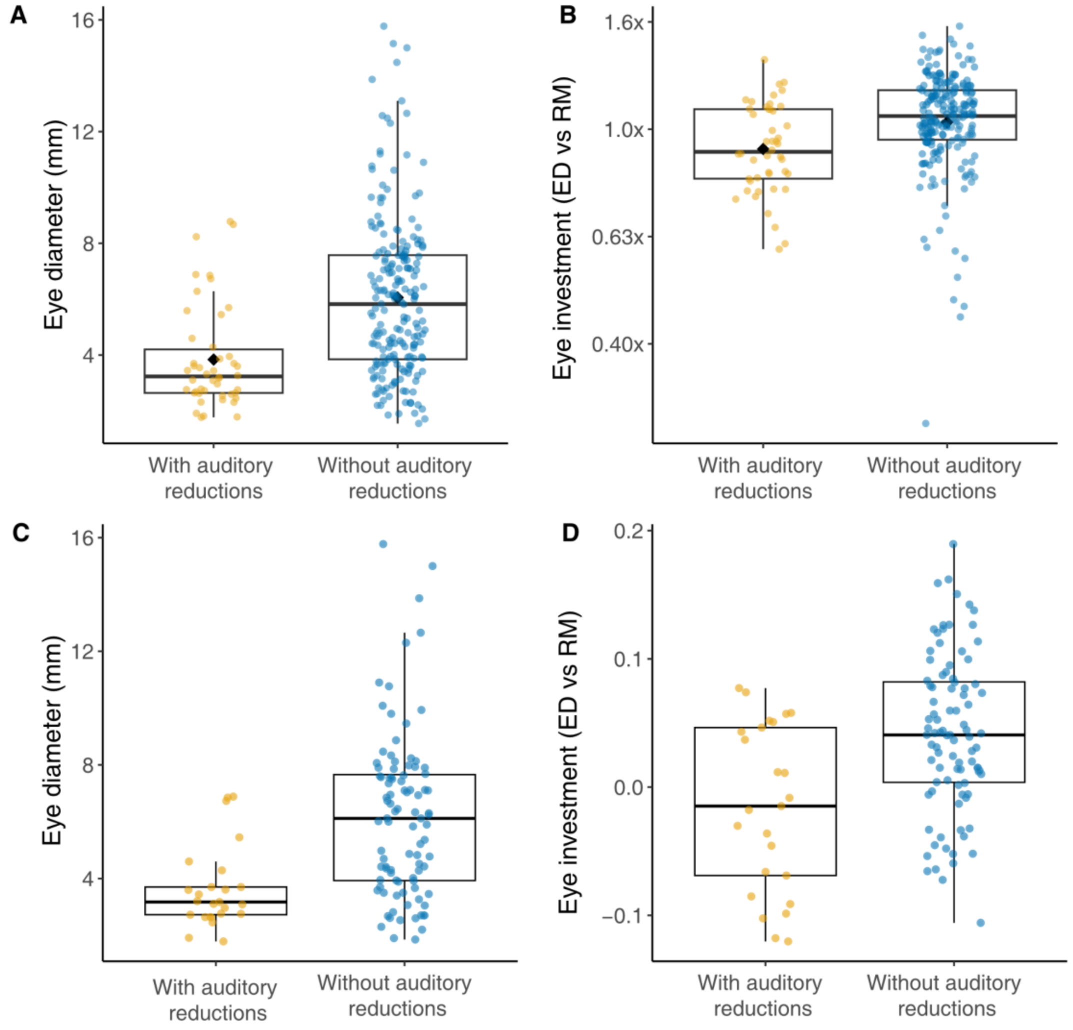


**Figure S3. Eye size differences between species with and without auditory reduction. (*A*)** Absolute eye diameter and **(*B*)** relative eye size (scaled by body mass) for species excluding fossorial taxa. **(*C*)** Absolute eye diameter and **(*D*)** relative eye size for ground-dwelling species only. Boxplots compare species with auditory reduction to those without. Across analyses, species with auditory loss exhibit significantly smaller eyes and reduced relative eye investment.

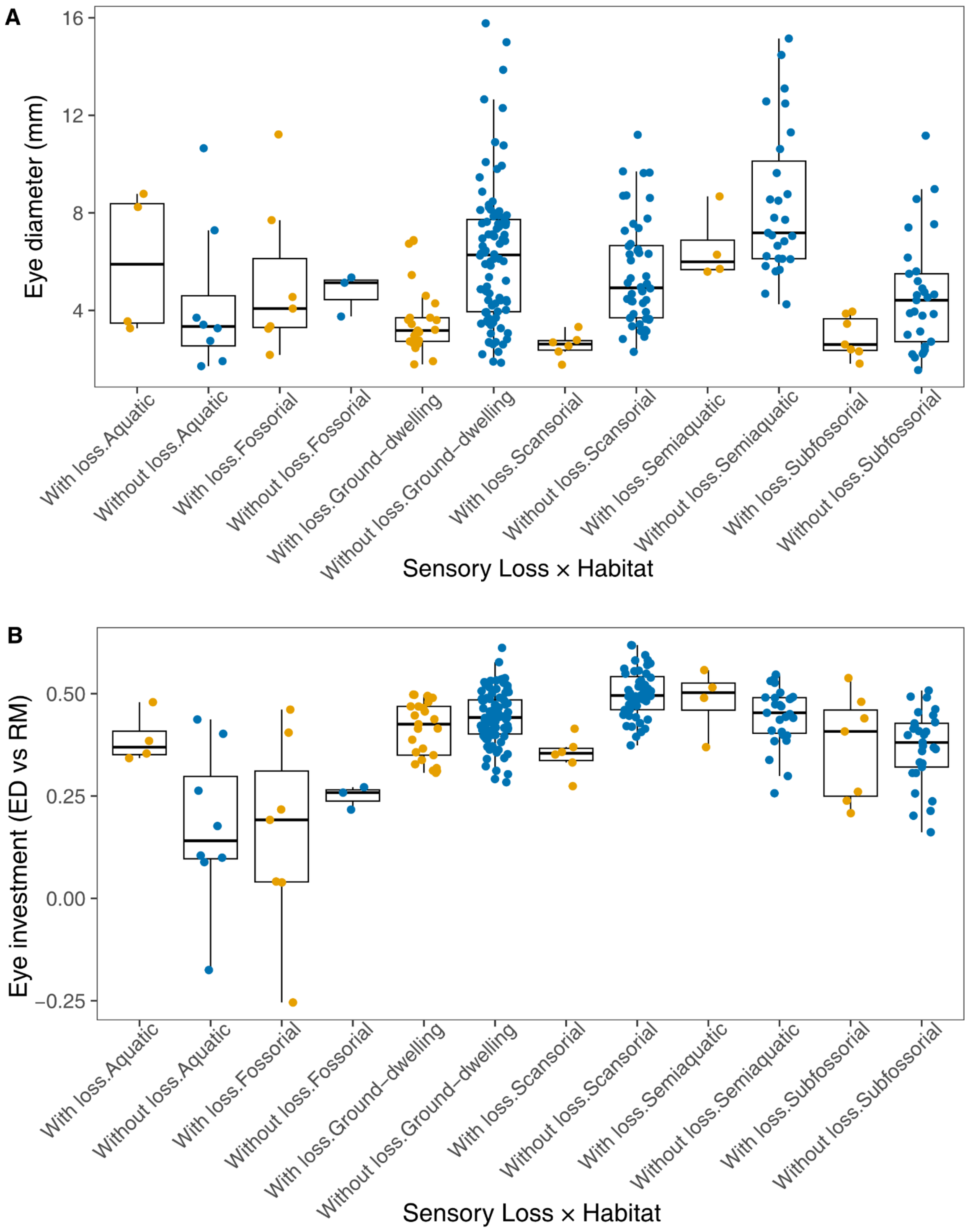


**Figure S4. Effects of auditory system loss and adult habitat on eye morphology. (*A*)** Absolute eye diameter and **(*B*)** relative eye size (scaled by body mass), derived from two-way phylogenetic ANCOVA analyses. Each boxplot displays the distribution of species values within each habitat category, separated by auditory system status. Auditory loss is associated with reduced eye size and investment across most habitats. All analyses include all species in the dataset.

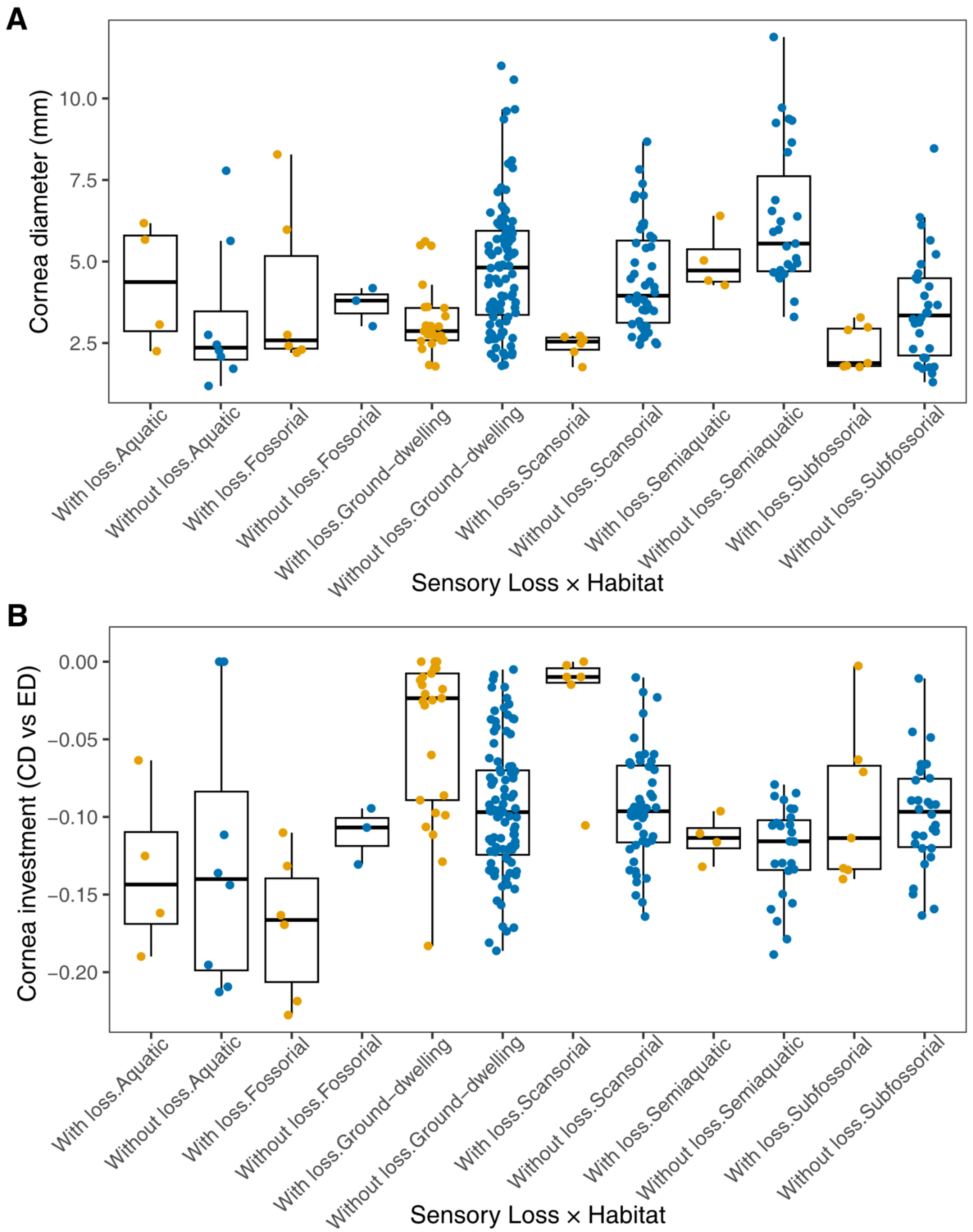


**Figure S5. Absolute and relative cornea size across habitats and auditory system conditions.** (***A***) Absolute cornea diameter and (***B***) relative corneal investment (CD *vs* ED), derived from two-way phylogenetic ANCOVA analyses. Each boxplot shows the distribution of species values within each habitat category, separated by auditory system status. Auditory loss is associated with reduced cornea size and slightly increased corneal investment across most habitats. All analyses include all species in the dataset.

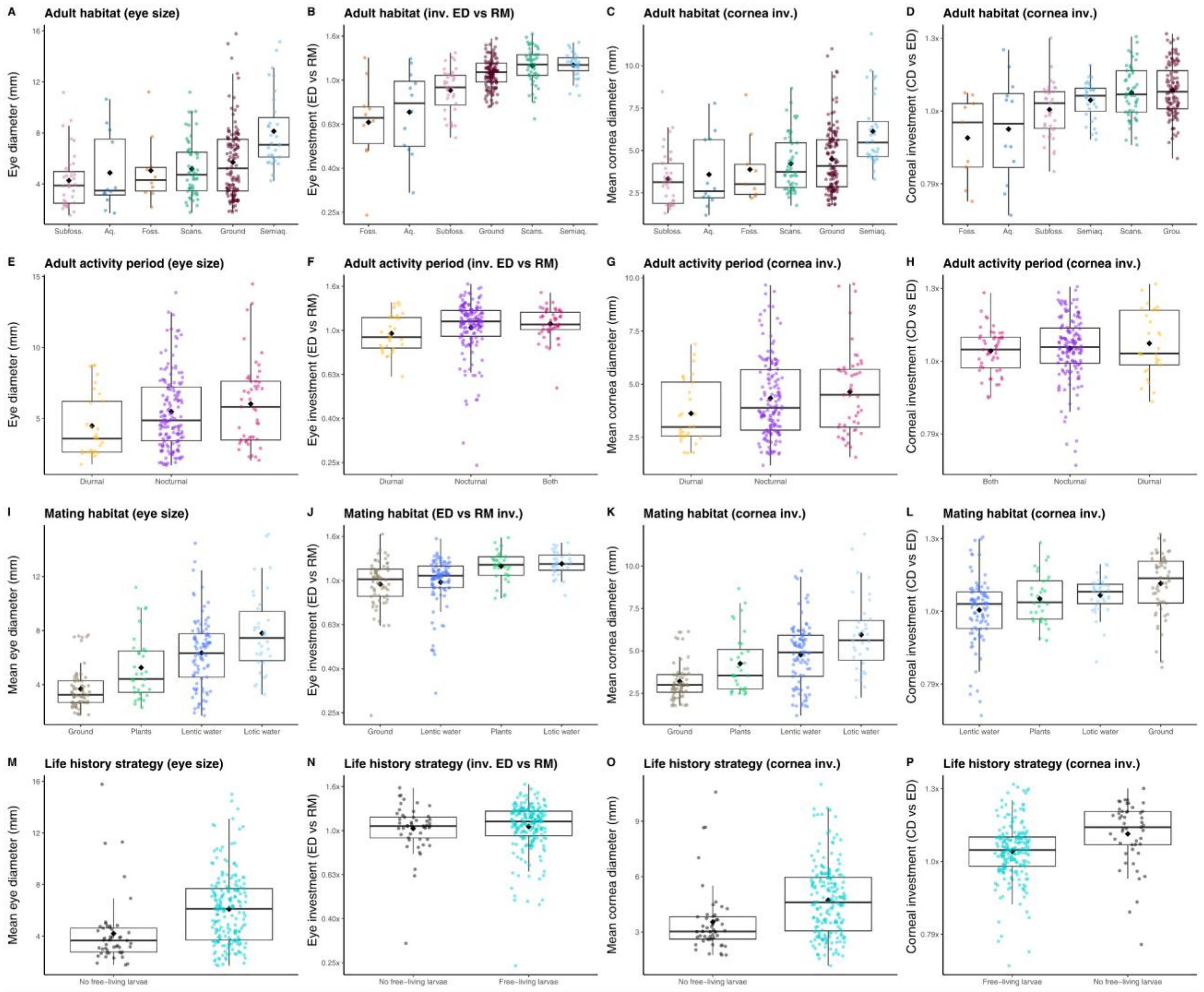


**Figure S6. Cornea and eye absolute size and investment differ significantly across ecological variables.** Utilizing different (A–D) adult habitats, (E–H) activity period, (I—L), mating habitat, and (M–P) life history strategy. All analyses include all species in the dataset.

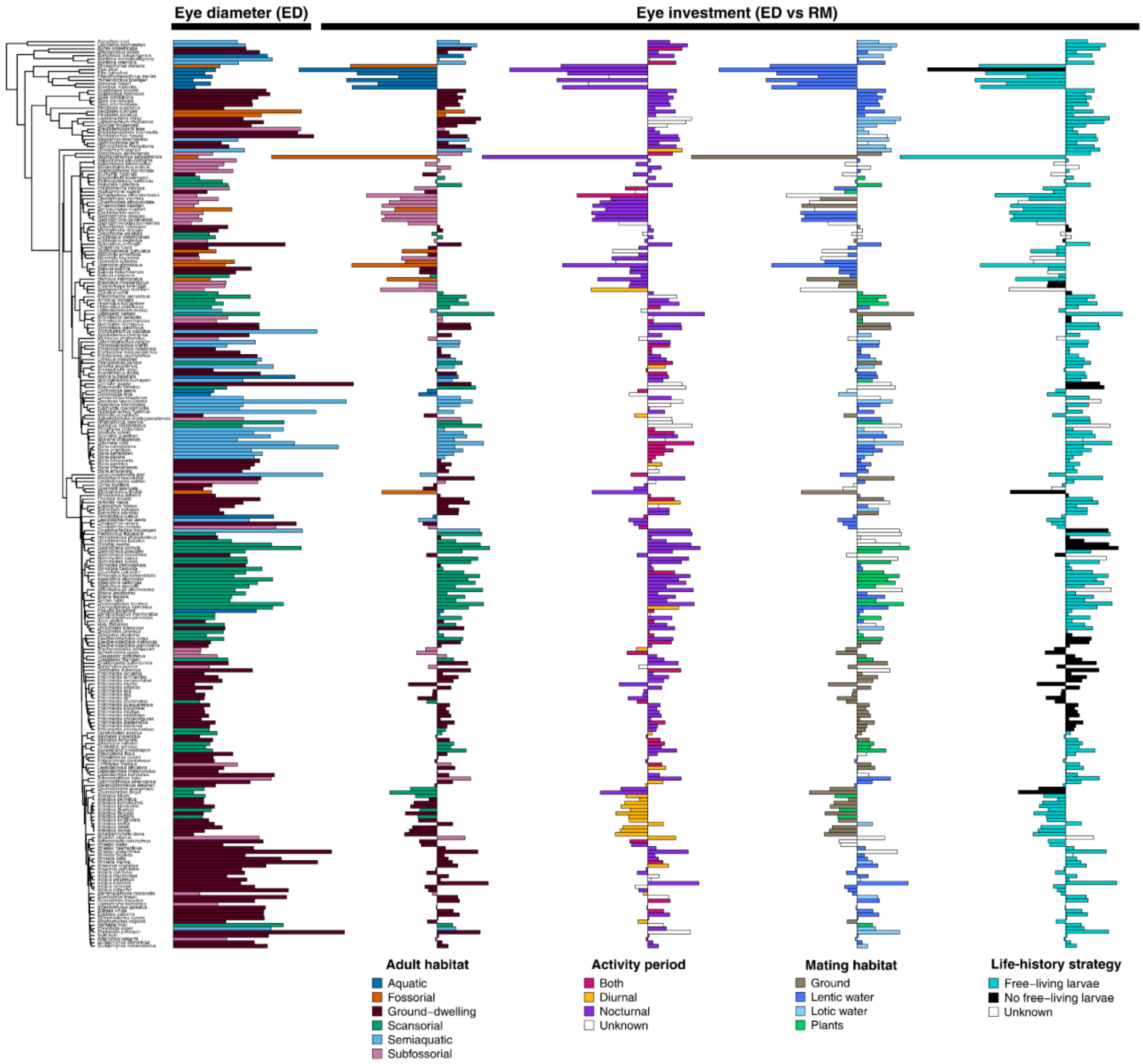


**Figure S7. Eye size and investment vary significantly across anurans with auditory and vocal reductions.** Phylogeny adapted from Portik et al. 2023 shows species means for absolute eye diameter (ED) and relative eye investment (ED x RM). All analyses include all species in the dataset.

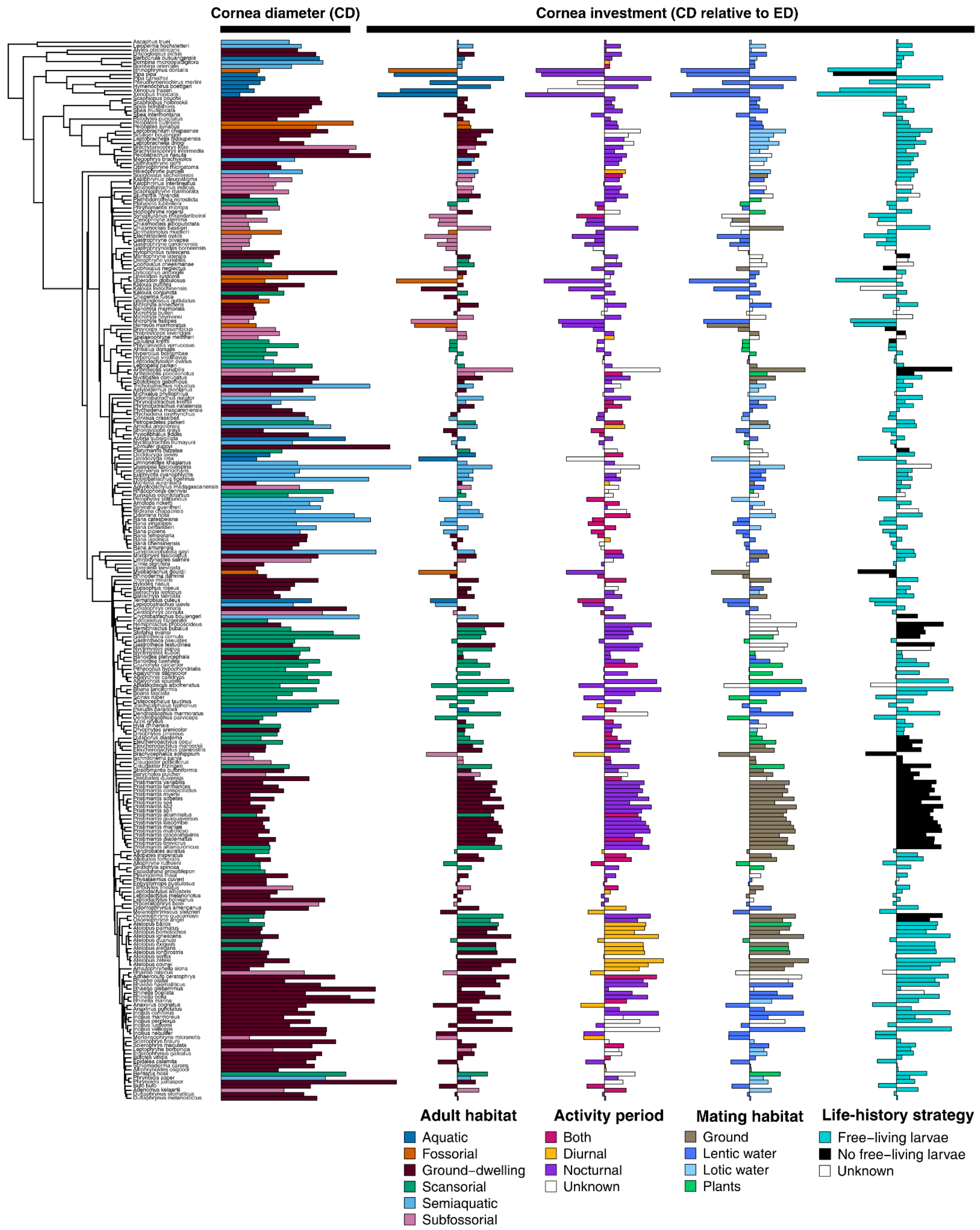


**Figure S8. Cornea size and investment vary significantly across anurans with auditory and vocal reductions.** Phylogeny adapted from Portik et al. 2023 shows species means for absolute cornea diameter (CD) and relative cornea investment (CD x ED). All analyses include all species in the dataset.
